# PELP1/SRC-3-dependent regulation of metabolic kinases drives therapy resistant ER+ breast cancer

**DOI:** 10.1101/2020.08.07.238550

**Authors:** Thu H. Truong, Elizabeth A. Benner, Kyla M. Hagen, Nuri A. Temiz, Carlos Perez Kerkvliet, Ying Wang, Emilio Cortes-Sanchez, Chieh-Hsiang Yang, Thomas Pengo, Katrin P. Guillen, Bryan E. Welm, Sucheta Telang, Carol A. Lange, Julie H. Ostrander

## Abstract

Recurrence of metastatic breast cancer stemming from acquired endocrine and chemotherapy resistance remains a health burden for women with luminal (ER+) breast cancer. Disseminated ER+ tumor cells can remain viable but quiescent for years to decades. Contributing factors to metastatic spread include the maintenance and expansion of breast cancer stem cells (CSCs). Breast CSCs frequently exist as a minority population in therapy resistant tumors. In this study, we show that cytoplasmic complexes composed of steroid receptor (SR) co-activators, PELP1 and SRC-3, modulate breast CSC expansion through upregulation of the HIF-activated metabolic target genes *PFKFB3* and *PFKFB4*. Seahorse metabolic assays demonstrated that cytoplasmic PELP1 influences cellular metabolism by increasing both glycolysis and mitochondrial respiration. PELP1 interacts with PFKFB3 and PFKFB4 proteins, and inhibition of PFKFB3 and PFKFB4 kinase activity blocks PELP1-induced tumorspheres and protein-protein interactions with SRC-3. PFKFB4 knockdown inhibited *in vivo* emergence of circulating tumor cell (CTC) populations in mammary intraductal (MIND) models. Application of PFKFB inhibitors in combination with ER targeted therapies blocked tumorsphere formation in multiple models of advanced breast cancer, including tamoxifen (TamR) and paclitaxel (TaxR) resistant models and ER+ patient-derived organoids (PDxO). Together, our data suggest that PELP1, SRC-3, and PFKFBs cooperate to drive ER+ tumor cell populations that include CSCs and CTCs.

**Significance:** Identifying non-ER pharmacological targets offers a useful approach to blocking metastatic escape from standard of care ER/estrogen (E2)-targeted strategies to overcome endocrine and chemotherapy resistance.

## INTRODUCTION

Metastatic recurrence is an incurable but common complication of ER+ breast cancer. Treatment of metastatic breast cancer typically results in endocrine resistance, and chemotherapy is largely ineffective in advanced disease. Altered signaling pathways drive therapy resistance and offer potential targets for metastatic ER+ breast cancer. PELP1 (proline, glutamic acid, leucine-rich protein 1) and SRC-3 (steroid receptor [SR] co-activator-3) have independently been shown to drive endocrine resistance. PELP1 and SRC-3 are both SR co-activators involved in normal development and cancer (1,2). Increased PELP1 expression is associated with higher tumor grade, tumor proliferation, and decreased breast cancer-specific survival (3,4). PELP1 is primarily nuclear in normal breast tissue; however, altered cytoplasmic PELP1 localization is observed in 40-58% of PELP1+ breast tumors (5). Analysis of breast tumor samples revealed that patients with high cytoplasmic PELP1 levels were less likely to respond to tamoxifen (tam) (4). Similarly, SRC-3 mRNA and protein overexpression are correlated with higher tumor grade and decreased overall and disease-free survival (6). SRC-3 overexpression is also linked to tam resistance in breast cancer models and human breast tumors (7,8). Both PELP1 and SRC-3 have essential nuclear functions, but also dynamically shuttle to the cytoplasm where they associate with signaling molecules and act as scaffolds for growth factor or SR pathways. These SR co-activators have emerged as promising targets in ER+ breast cancer and as potential mediators of therapy resistance.

Cancer stem cells (CSCs) are poorly proliferative and frequently exist as a minority sub-population of cells that drive therapy resistance and metastasis (9). In contrast to non-CSCs, breast CSCs form colonies in serum-free suspension culture (i.e. tumorspheres), express stem cell markers (e.g. ALDH+ or CD44^hi^/CD24^lo^), exhibit enhanced resistance to chemo and endocrine therapies, and express markers of epithelial to mesenchymal transition (EMT). The ability to survive and self-renew following treatment allows CSCs to evade standard chemo and endocrine therapies aimed at rapidly dividing cancer cells and to drive metastatic tumor growth.

Growing evidence has implicated SR co-activators as mediators of CSC self-renewal. For example, SRC-3 drives CSC formation and tumor outgrowth in breast cancer models. Treatment with SI-2, an SRC-3 inhibitor, decreased SRC-3-induced CSC formation in breast cancer cell and xenograft models (10). Our laboratory reported that cytoplasmic complexes composed of PELP1 and SRC-3 mediate breast CSC expansion (11). Targeting SRC-3 using shRNA or pharmacological inhibitors (i.e. SI-2) abrogated PELP1/SRC-3 complex formation, PELP1-induced tumorspheres, and expression of PELP1 target genes that promote cancer cell survival. These studies imply that inhibiting PELP1 and its binding partners may provide a way to target the breast CSC population in order to improve patient outcomes.

Herein we sought to identify the molecular mechanisms that contribute to PELP1-driven CSC survival and self-renewal in ER+ breast cancer. Using endocrine and chemotherapy resistant breast cancer models, our findings suggest that PELP1/SRC-3 complexes modulate the CSC compartment through gene programs associated with metabolic adaptation. In contrast to current therapies that fail to adequately target slow-growing breast CSCs, our studies reveal therapy combinations that inhibit cooperating signaling cascades, while simultaneously targeting ER. By targeting CSCs directly, this approach promises to significantly improve the lives of patients with recurrent ER+ breast cancer.

## MATERIALS AND METHODS

### Cell Culture

STR authentication was performed by ATCC (October 2018). MCF-7 PELP1 and J110 cells were cultured as described (11). MCF-7 (12) and T47D TamR (13) cells were cultured in 100 nM tamoxifen. MCF-7 TaxR (14) cells were cultured in 2 μM Taxol. For 3D (tumorsphere) conditions, cells were cultured as described (11).

### Patient-Derived Organoids (PDxO)

PDxOs (HCI-003, −011, −017) were cultured in Advanced DMEM/F12 (Thermo Fisher) containing 5% FBS, 1X HEPES, 1X GlutaMax, 50 μg/ml Gentamicin (Genesee), 1 μg/ml hydrocortisone, 10 ng/ml EGF, and supplemented with 10 μM Y-27632 (Selleckchem), 100 ng/ml FGF2 (PeproTech), and 1 mM N-acetyl cysteine (Sigma). PDxOs were embedded into Matrigel (growth factor reduced; Corning) using the hanging drop method into 6-well plates and passaged every ~14-18 days.

### Synergy Screens

PDxOs were grown as described above in 384-well plates: Prior to assay, PDxOs were cultured in PDxO media containing charcoal stripped FBS and allowed to incubate overnight. PDxOs were disbursed with Dispase and re-seeded in PDxO media on 384-well plates coated with Matrigel. Following 24h after seeding, plates were dosed with SI-2 (0-2 μM), 5MPN (0-50 μM), and tam (0-10 μM) using a robotic assisted pipette (ViaFlo). On Day 6 post-treatment, a Cell Titer Glo 3D assay (Promega) was performed per manufacturer’s instructions. Luminescence readings were measured using an Envision Spectrophotometer. Fold changes were calculated using Day 0 baseline readings. % survival was calculated for each well, and ZIP synergy model (15) were run to determine synergy scores with the R package synergy finder (16). Independent drug response was calculated using the four-parameter logistic model (4-PL). 3 biological replicates, each one consisting of 4 technical replicates per dosing, was performed.

### Mammary Intraductal (MIND) Model

Intraductal injections of single cells were performed as described (17,18). Seven-week old female NSG mice (#005557) were purchased from Jackson Laboratory. Five mice/group were injected with 5 x 10^4^ cells into each nipple of the 4^th^ inguinal glands with the indicated breast cancer cell line. Mammary glands were harvested 8 weeks after injection, fixed in 4% PFA, and processed for H&E staining. H&E sections were analyzed using ImageJ or Q-path to quantitate the total mammary gland area (%) that contained tumor cells.

### Statistical Analysis

Data were tested for normal distribution using Shapiro-Wilks normality test and homogeneity of variances using Bartlett’s Test. Once data met these two requirements, statistical analyses were performed using one-way or two-way ANOVA in conjunction with Tukey multiple comparison test for means between more than two groups or Student *t* test for means between two groups, where significance was determined with 95% confidence. For the MIND study with four groups defined by two factors (cyto PELP1 vs. WT PELP1, and shPFKFB4 vs. shGFP), a regression model identified a significant interaction due to shPFKFB4 at an alpha level of 0.1 (p=0.084).

## RESULTS

### Cytoplasmic PELP1 promotes CSCs and HIF-regulated gene expression

Breast CSCs represent a minority of the total cell population (1-5%) (19), making it difficult to detect CSC-specific changes in bulk tumor populations. We therefore measured breast CSC frequency by comparing ALDH activity (**Figure 1A**, **Supplementary Figure 1**) and CD44^hi^/CD24^lo^ ratios (**Figure 1B**, **Supplementary Figure 2**) in MCF-7 cells stably expressing LXSN (vector control), WT PELP1, or cytoplasmic (cyto) PELP1 cultured in either 2D (adherent) or 3D (tumorsphere) conditions. Relative to 2D, 3D conditions increased breast CSC markers in MCF-7 cells expressing LXSN, WT PELP1, or cyto PELP1 (**Figure 1A, 1B**). In 2D conditions, cyto PELP1 expressing cells had no significant changes in ALDH activity when compared to LXSN or WT PELP1; however, when the same models were cultured in 3D conditions, ALDH activity was significantly increased in cells expressing cyto PELP1 (12.0% ± 2.9) compared to LXSN (6.6% ± 0.67, p = 0.023) and WT PELP1 (2.6% ± 0.76, p = 0.0015). In 2D conditions, CD44^hi^/CD24^lo^ populations were increased in cyto PELP1 expressing cells (13.0% ± 0.49) compared to LXSN (2.6% ± 0.042, p < 0.0001) or WT PELP1 (1.2% ± 0.19, p < 0.0001), and this trend was enhanced in 3D conditions (cyto PELP1, 19.4% ± 1.4; LXSN, 9.0% ± 1.1, p = 0.0045; WT PELP1, 2.3% ± 0.18, p = 0.0011). WT PELP1 displayed lower ALDH activity and CD44^hi^/CD24^lo^ ratios relative to LXSN controls, suggesting that nuclear PELP1 limits CSC behavior. These results indicate that both 3D culture and cyto PELP1 expression independently increase CSC expansion in MCF-7 cell models.

**Figure 1.**
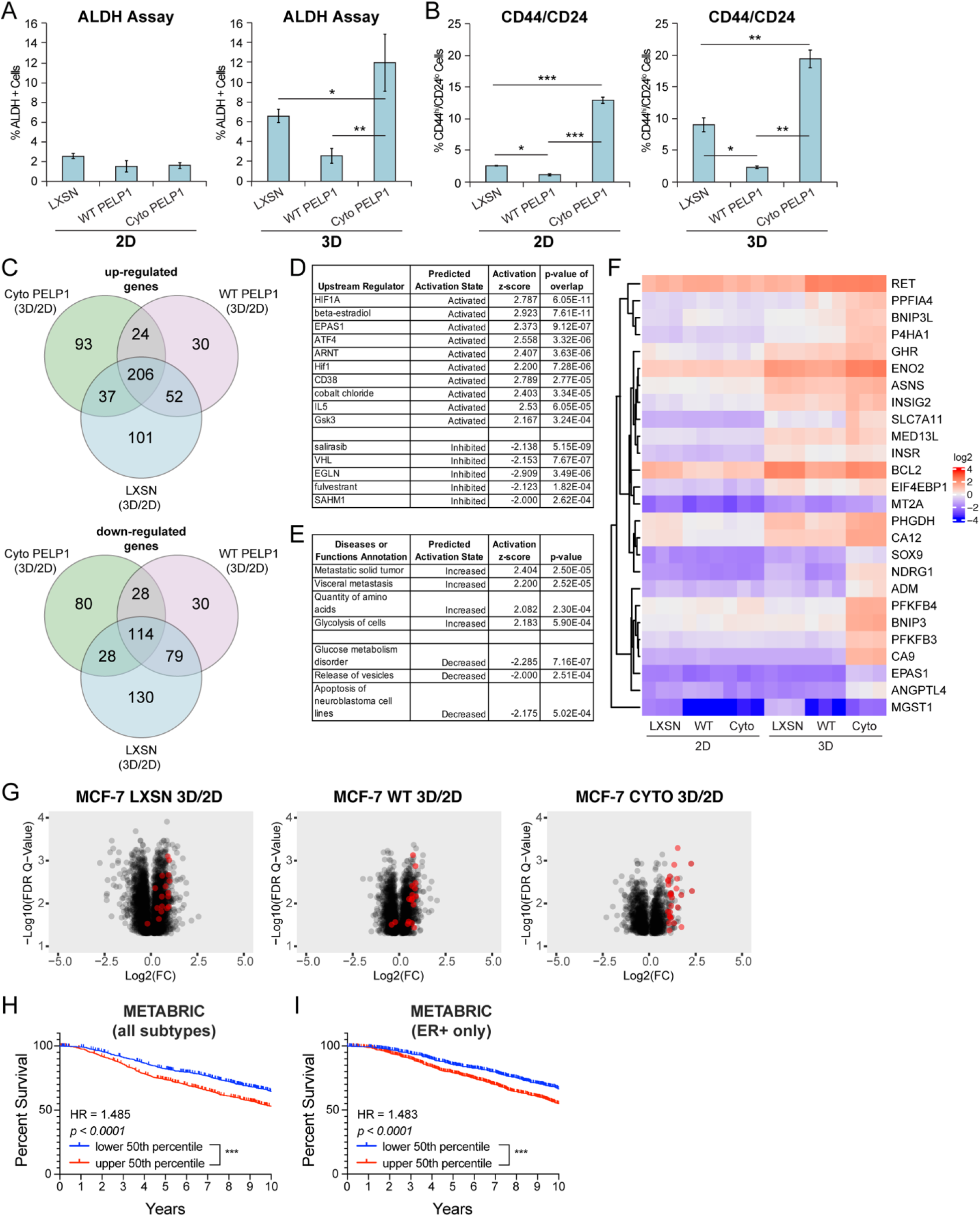
PELP1-induced gene expression is altered in 3D conditions. (**A**) ALDH activity and (**B**) CD44^hi^/CD24^lo^ populations in MCF-7 PELP1 cells. (**C**) Venn diagrams showing unique genes up or downregulated >2-fold in MCF-7 PELP1 cells (3D vs. 2D). IPA analysis of (**D**) upstream regulators and (**E**) diseases or functions. (**F**) Representative heat-map showing log2(FPKM) values of cyto PELP1 gene signature. (**G**) Volcano plots of 3D vs. 2D comparison of MCF-7 PELP1 cells. X-axis is Log2(fold change) and Y-axis represent −Log 10 Benjamini-Hochberg corrected Q-values. Kaplan-Meier curves for upper and lower 50^th^ percentile of cyto PELP1 gene signature expression in the METABRIC (**H**) all subtypes and (**I**) ER+ only patient cohorts. Graphed data represent the mean ± SD (n = 3). * p < 0.05, ** p < 0.01, *** p < 0.001.

We performed RNA-seq on MCF-7 PELP1 models grown as 3D tumorspheres and compared these data to studies conducted in 2D culture (11) to identify candidate genes and pathways differentially regulated in cyto PELP1 expressing cells. Comparison of 3D versus 2D conditions identified 206 upregulated and 114 downregulated genes similarly regulated by >2-fold in all cell lines (LXSN, WT PELP1, cyto PELP1) (**Figure 1C, Supplementary Figure 3**). Ingenuity Pathway Analysis (IPA) of these 320 genes revealed activation of estrogen, growth factor, cytokine, and NF-κB pathways (**Supplementary Table 1**). Significantly activated and inhibited “Diseases and Functions” are summarized in **Supplementary Table 2**. 3D to 2D comparison in cyto PELP1 expressing cells identified 173 differentially expressed genes (93 upregulated, 80 downregulated) compared to LXSN or WT PELP1 (**Figure 1C, Supplementary Figure 4**). These 173 genes were analyzed with IPA to identify cyto PELP1-specific pathways (**Figure 1D**), biological functions, or disease states (**Figure 1E**), and predicted increased HIF activation, estradiol, ATF4, and glycolytic-mediated pathways. We created representative heatmaps to illustrate 3D-specific regulation in upstream regulator analysis associated with HIF and ATF4 pathway activation (>2-fold; **Figure 1F**) and generated a cyto PELP1 upregulated gene signature (**Supplementary Table 3**). Volcano plots of differentially regulated genes are shown in **Figure 1G**; red dots indicate genes in the cyto PELP1 signature. We then used the cyto PELP1 upregulated gene signature to query the METABRIC breast cancer database. Higher expression of this gene signature was associated with lower overall survival (OS) in the METABRIC cohort (hazard ratio = 1.485, p < 0.0001, **Figure 1H**). We tested this on the ER+ only subtype within the METABRIC cohort and found similar results (hazard ratio = 1.483, p < 0.0001, **Figure 1I**). A similar query of the TCGA database revealed no significant differences in OS (**Supplementary Figure 5**). Taken together, these data identify genes involved in cyto PELP1-mediated pathways that promote CSCs, including those associated with HIF-activated and glycolytic pathways.

### Cytoplasmic PELP1 drives metabolic plasticity

Given the strong activation of HIF and metabolic pathways detected in the RNA-seq analysis, we used qPCR to test HIF-activated target genes. HIF activates the *PFKFB* family, which are metabolic bi-functional kinase/phosphatases (20). We found that mRNA levels of *EPAS1* (i.e. HIF2α), *PFKFB3*, and *PFKFB4* were upregulated in cells expressing cyto PELP1 relative to LXSN or WT PELP1 in 3D, but not 2D conditions (**Figure 2A**). Additional validation of HIF-activated metabolic and stem cell genes include *NDRG1* and *SOX9* (**Figure 2A**). Given the central role of HIF pathways in metabolism (21), we investigated the effect of PELP1 on metabolic pathways using the Seahorse Cell Energy Phenotype test to measure oxygen consumption rate (OCR) and extracellular acidification rate (ECAR). At baseline, MCF-7 cells expressing cyto PELP1 exhibited a significant increase in OCR levels compared to LXSN and WT PELP1. Under stressed conditions (i.e. after FCCP and oligomycin), OCR was increased in cyto PELP1 expressing cells compared to LXSN (p = 0.0096). ECAR was significantly different in cyto PELP1 expressing cells compared to LXSN at baseline, but WT and cyto PELP1 displayed an increase in ECAR compared to LXSN controls (p = 0.046 and 0.0045) under stressed conditions (**Figure 2B**). To systematically test effects on key parameters of mitochondrial function, we performed the Seahorse Mito Stress test. Cyto PELP1 expression significantly increased basal respiration, compared to LXSN and WT PELP1 (p < 0.0001 and 0.0001). Furthermore, cyto PELP1 increased ATP-linked respiration, proton leak, maximal respiration, and non-mitochondrial respiration (**Figure 2C**). Cyto PELP1 expressing cells had a 4-fold increase in glucose uptake compared to WT PELP1 and LXSN, as measured by 2-NBDG (**Figure 2D, Supplementary Figure 6**). Collectively, these results indicate cyto PELP1 drives HIF-activated metabolic programs (i.e. *PFKFB3, PFKFB4*) in 3D culture, and affects mitochondrial respiration and glycolysis, indicative of metabolic plasticity.

**Figure 2.**
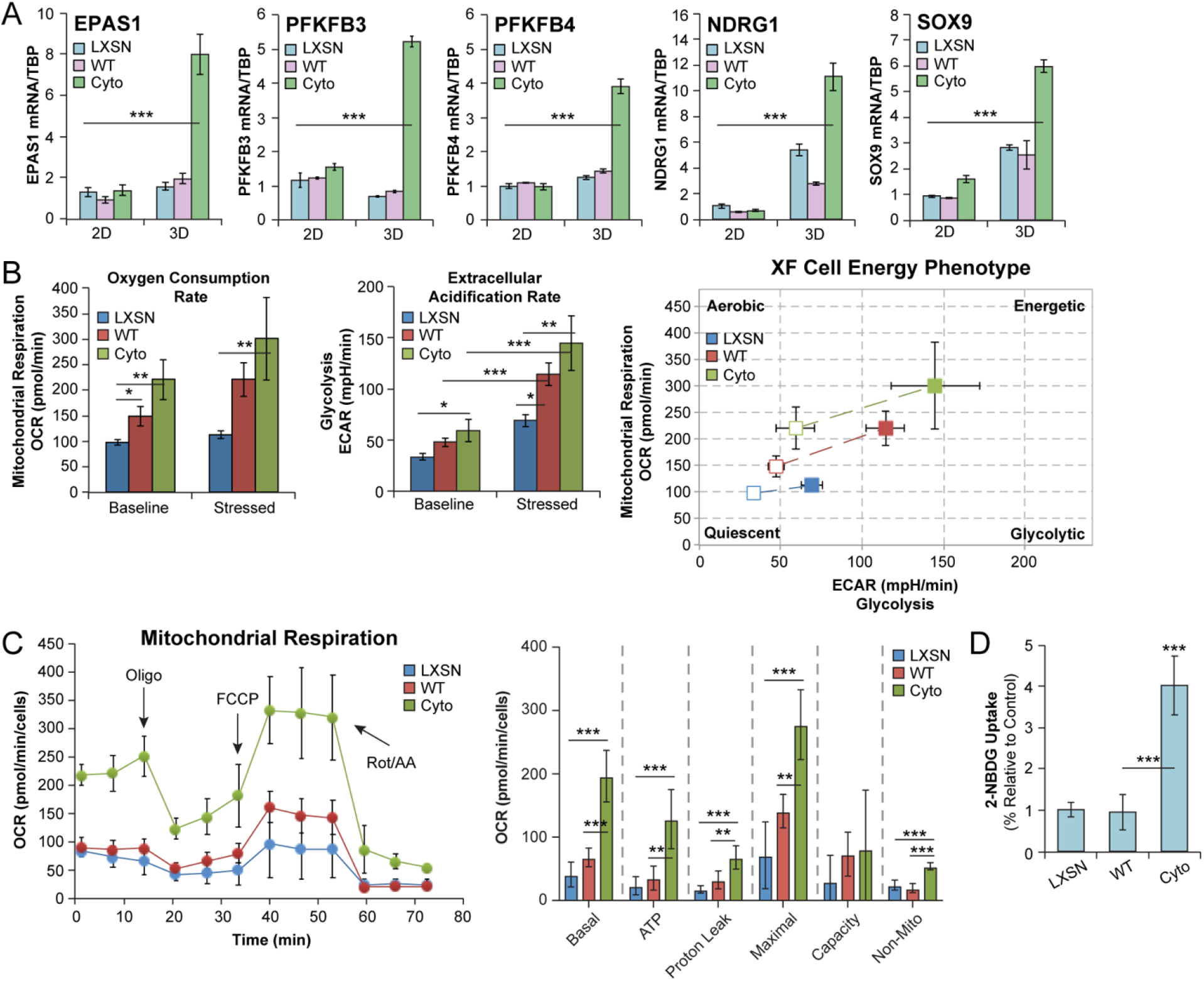
PELP1 cytoplasmic signaling upregulates HIF-activated metabolic pathways. (**A**) mRNA levels of *EPAS1, PFKFB3, PFKFB4, NDRG1*, and *SOX9* in MCF-7 PELP1 cells. (**B**) OCR and ECAR measured in MCF-7 PELP1 cells by Seahorse Cell Energy Phenotype test. (**C**) OCR measured in MCF-7 PELP1 cells by Seahorse Mito Stress test. (**D**) Glucose uptake in cells treated with 2-NBDG (10 μM). 2-NBDG uptake is represented as % cells relative to control. Graphed data represent the mean ± SD (n = 3). * p < 0.05, ** p < 0.01, *** p < 0.001.

### Inhibition of PFKFBs disrupts PELP1/SRC-3 complexes and tumorsphere formation

We hypothesized that HIF-activated targets PFKFB3 and PFKFB4 are required components of the PELP1/SRC-3 complex. Co-immunoprecipitation of PFKFB3 or PFKFB4 demonstrated increased association with PELP1 in cells expressing cyto PELP1 relative to LXSN or WT PELP1 (**Figure 3A, 3B**). Treatment with PFK158 and 5MPN, inhibitors of PFKFB3 and PFKFB4 respectively, reduced the PELP1/SRC-3 interaction (**Figure 3C, 3D**). These inhibitors also blocked PELP1/PFKFB3 and PELP1/PFKFB4 (**Supplementary Figure 7A, 7B**) interactions in cyto PELP1 expressing cells; similar results were observed with another PFKFB3 inhibitor (PFK15; **Supplementary Figure 7C, 7D**).

**Figure 3.**
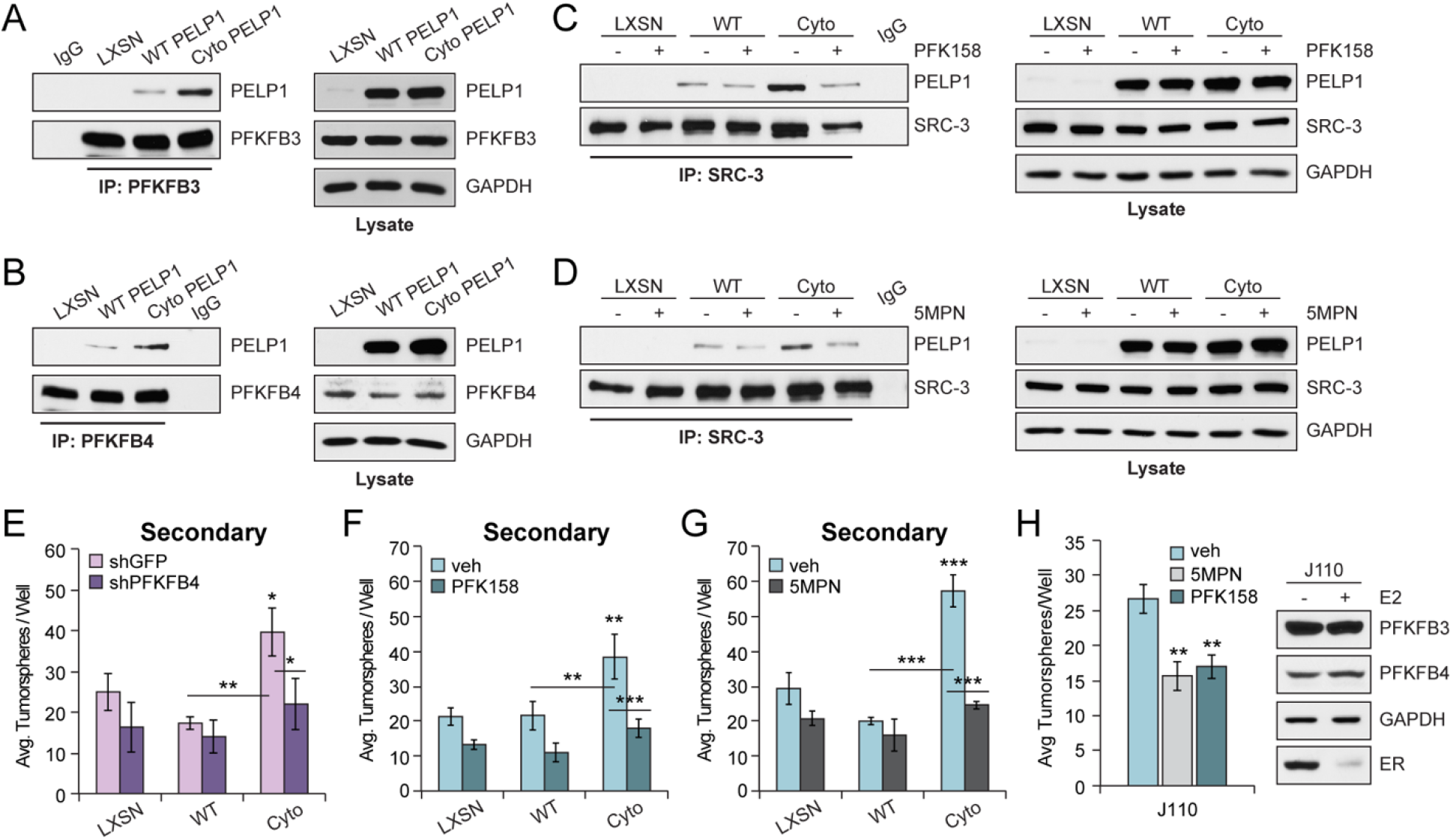
PFKFB inhibition blocks PELP1/SRC-3 signaling. Co-immunoprecipitation of (**A**) PELP1 and PFKFB3 or (**B**) PFKFB4 in MCF-7 PELP1 cells. Co-immunoprecipitation of PELP1 and SRC-3 in MCF-7 PELP1 cells treated with vehicle (DMSO), (**C**) PFK158 (100 nM), or (**D**) 5MPN (5 μM). Cell lysate controls (*right*). (**E**) Secondary tumorsphere assays in MCF-7 PELP1 shGFP control or shPFKFB4 knockdown cells. Secondary tumorsphere assays in MCF-7 PELP1 cells treated with vehicle, (**F**) PFK158 or (**G**) 5MPN. (**H**) Secondary tumorsphere assays in J110 cells treated with vehicle, PFK158, or 5MPN. Western blot shows PFKFB3 and PFKFB4 protein in J110 cells. Graphed data represent the mean ± SD (n = 3). * p < 0.05, ** p < 0.01, *** p < 0.001.

Next, we tested the effect of PFKFB inhibition on cyto PELP1-induced tumorspheres, an *in vitro* assay to assess breast CSC activity (11). PFKFB4 knockdown (**Supplementary Figure 8**) decreased tumorsphere formation in cyto PELP1 expressing cells by ~50%, but not in LXSN or WT PELP1 (**Figure 3E**, p = 0.0103). Attempts to stably knockdown PFKFB3 were not successful, suggesting that PFKFB3 is crucial for cell viability (22). Inhibitors of PFKFB3 and PFKFB4 reduced cyto PELP1-induced tumorspheres, but had no effect on cells expressing either LXSN or WT PELP1. (**Figure 3F, 3G; Supplementary Figure 7E**). To evaluate PFKFB inhibitors in an alternative PELP1/SRC-3 model, we used a murine tumor cell line (J110) established from the MMTV-SRC-3 mouse (23). Similar to MCF-7 PELP1 models, PFK158 or 5MPN inhibited tumorsphere formation by ~40% in J110 cells (**Figure 3H**). Western blotting indicated that PFKFB3 and PFKFB4 protein levels remained unchanged in response to E2, while ER levels decreased, presumably due to ligand-induced turnover (**Figure 3H**, right). These results indicate that blocking PFKFB3 or PFKFB4 through knockdown or pharmacological inhibition disrupts expansion and self-renewal of PELP1-driven CSC populations.

### PFKFB4 reduces *in vivo* expansion of CTCs in cyto PELP1 MIND xenografts

To evaluate if PELP1 promotes tumor formation *in vivo*, we injected MCF-7 WT and cyto PELP1 expressing cells (5 x 10^4^) into the inguinal mammary glands of adult female mice (6-8 week old, 4 mice/group) to generate mammary intraductal (MIND) tumors. Both cell lines had 100% engraftment rates (**Figure 4A, Supplementary Figure 9**). Tumor area (%) calculated from H&E images of each mammary gland revealed increased tumor volume in cyto PELP1 (25.7% ± 16.5) compared to WT PELP1 MIND xenografts (10.9% ± 9.5, p = 0.046) (**Figure 4B**).

**Figure 4.**
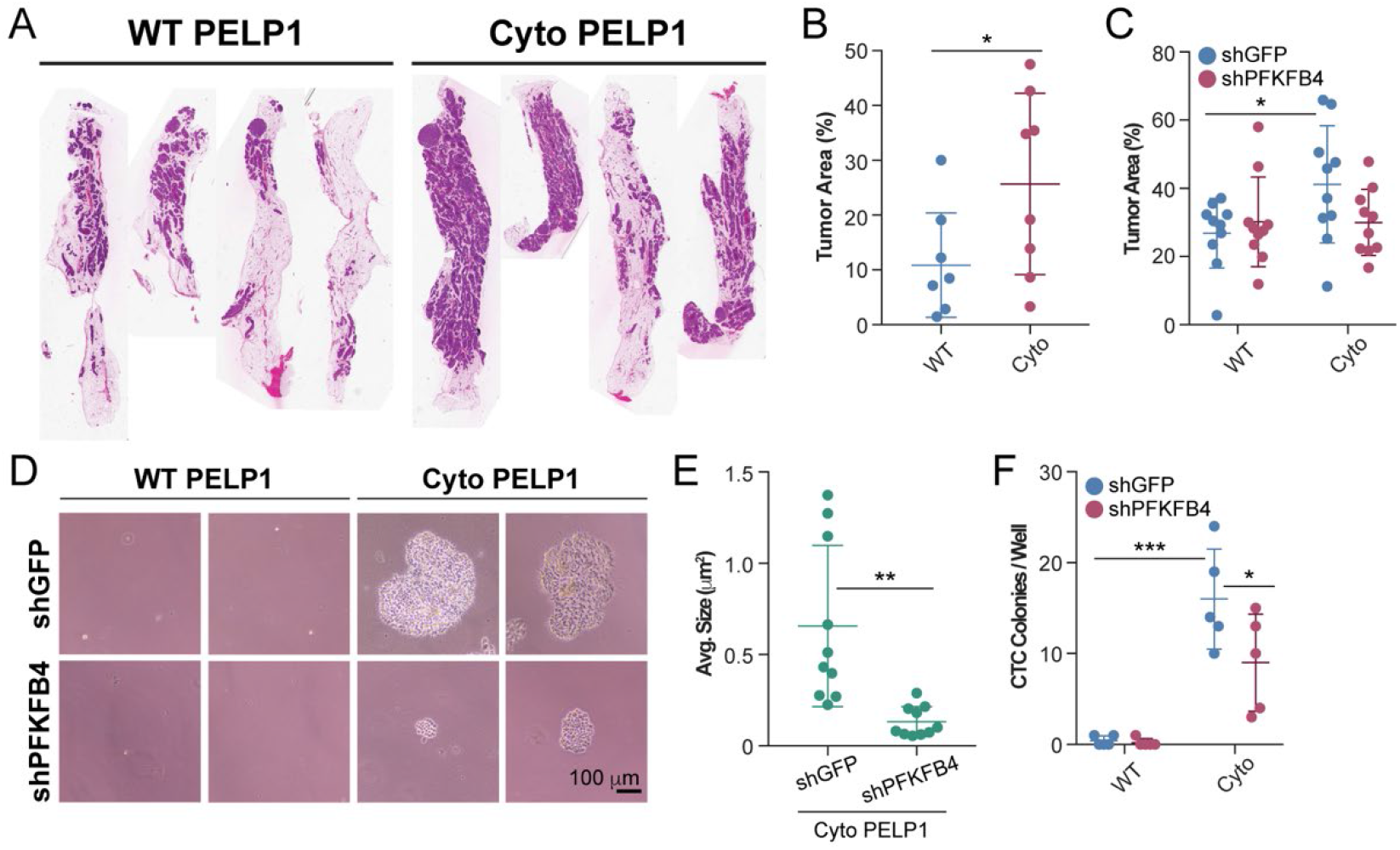
PFKFB4 knockdown abrogates cyto PELP1 CTCs in MIND xenograft models. (**A**) Representative H&E stains from MIND glands (WT and cyto PELP1). (**B**) Tumor area (%) calculated from H&E sections from (**A**). (**C**) Tumor area (%) calculated from H&E sections from WT and cyto PELP1 (shGFP, shPFKFB4) MIND glands. (**D**) Representative images of CTCs from blood samples collected from mice injected with WT or cyto PELP1 (shGFP, shPFKFB4) cells. (**E**) Average size of soft agar colonies (CTCs) from (**D**). (**F**) Average number of colonies/well (CTCs). Graphed data represent the mean ± SD (n = 5). * p < 0.05, ** p < 0.01, *** p < 0.001.

Based on our *in vitro* data showing that inhibition of PFKFB4 (knockdown and 5MPN) decreased tumorspheres, we queried PFKFB4 mRNA levels on OS in METABRIC datasets. High PFKFB4 mRNA expression is associated with decreased OS in all subtypes and ER+ only patient cohorts (**Supplementary Figure 10**). Therefore, we tested whether PFKFB4 knockdown would impact MIND tumor growth or the presence of circulating tumor cells (CTCs); a marker of metastatic potential and associated CSC behavior (24). 5 mice/group were injected with MCF-7 WT or cyto PELP1 expressing cells harboring either shGFP control or shPFKFB4. 8 weeks post-injection, mammary glands were fixed and processed for H&E staining (**Supplementary Figure 11**). As in **Figure 4B**, the difference in means between WT PELP1 shGFP (26.8% ± 10.2) and cyto PELP1 shGFP (41.2% ± 17.2) tumor area remained significant (p = 0.036, **Figure 4C**). However, knockdown of PFKFB4 in MCF-7 cells expressing either WT PELP1 or cyto PELP1 failed to significantly affect primary tumor growth. To assess disseminated tumor cells, blood samples were collected during euthanization and seeded into soft agar assays to detect CTCs. Mice injected with WT PELP1 (shGFP or shPFKFB4) expressing cells did not exhibit CTC colony formation. In sharp contrast, blood samples from mice engrafted with cyto PELP1 cells developed large viable colonies, indicating the presence of CTCs. Knockdown of shPFKFB4 in MCF-7 cyto PELP1 expressing cells reduced both colony formation (p < 0.0492) and colony size (p < 0.0016) (**Figure 4D-4F**). These data demonstrate a requirement for PFKFB4 in cyto PELP1-driven CTC formation and expansion *in vivo*.

### Targeting PELP1/SRC-3 complexes in therapy resistant breast cancer and PDxO models

Paclitaxel (Taxol) is a chemotherapy used to treat late stage breast cancer. Increased PELP1, HIF1α, and HIF-2α expression has been observed in triple negative breast cancer (TNBC) cells in response to Taxol (14). To evaluate whether PELP1 expression affects response to Taxol in ER+ breast cancer, we treated MCF-7 PELP1 cells (LXSN, WT PELP1, cyto PELP1) cultured as tumorspheres with Taxol (0 to 125 nM). We assessed tumorsphere formation and calculated IC50 values for each cell line (**Figure 5A**). IC50 (Taxol) for cyto PELP1 expressing cells was ~2-fold higher than LXSN or WT PELP1. These results suggest that cyto PELP1 expression confers enhanced Taxol resistance compared to LXSN or WT PELP1.

**Figure 5.**
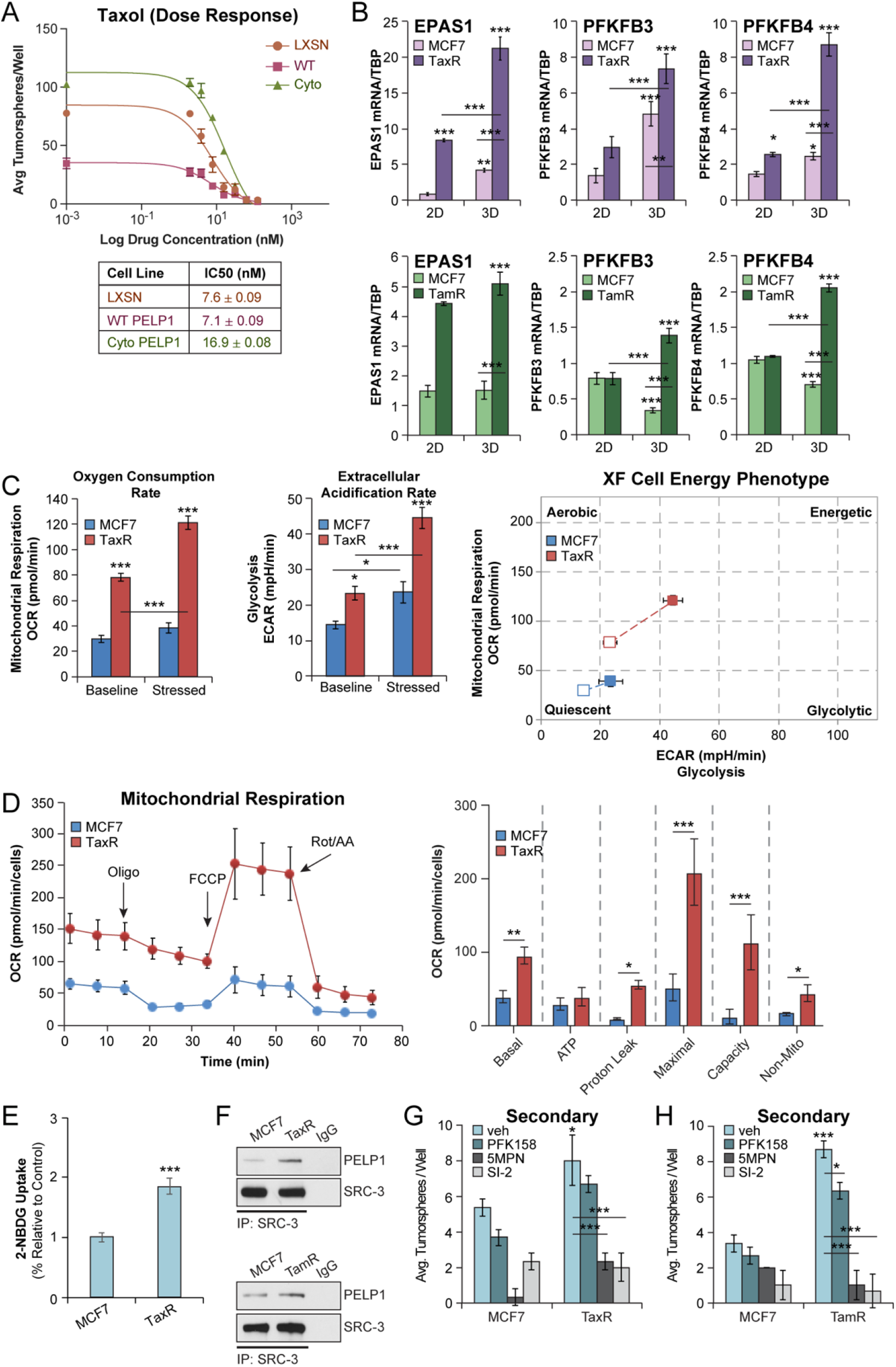
Therapy resistant models phenocopy cyto PELP1 cancer biology. (**A**) Taxol dose response in MCF-7 PELP1 cells (0-125 nM Taxol). (**B**) mRNA levels of *EPAS1, PFKFB3*, and *PFKFB4* in MCF-7 TaxR (*top*) or TamR (*bottom*) cells cultured in 2D or 3D conditions. (**C**) OCR and ECAR measured in MCF-7 TaxR cells by Seahorse Cell Energy Phenotype test. (**D**) OCR measured in MCF-7 TaxR cells by Seahorse Mito Stress test. (**E**) Glucose uptake in cells treated with 2-NBDG (10 μM). (**F**) Co-immunoprecipitation of PELP1 and SRC-3 in MCF-7 TaxR (*top*) or TamR (*bottom*) cells. Secondary tumorsphere assays in (**G**) MCF-7 TaxR and (**H**) MCF-7 TamR cells treated with vehicle (DMSO), PFK158 (100 nM), 5MPN (5 μM), or SI-2 (100 nM). Graphed data represent the mean ± SD (n = 3). * p < 0.05, ** p < 0.01, *** p < 0.001.

Next, we determined if PELP1/SRC-3 signaling mediates therapy resistance in tamoxifen resistant (TamR) and paclitaxel-resistant (TaxR) cell lines. HIF and cyto PELP1 regulated genes, *EPAS1, PFKFB3, and PFKFB4*, mRNA levels were increased in MCF-7 TaxR (**Figure 5B**, *top*) and TamR cells (**Figure 5B**, *bottom*) relative to MCF-7 parental controls, particularly in 3D conditions. 3D PELP1 target genes, *NDRG1* and *SOX9* were also upregulated in TaxR and TamR cells relative to parental MCF-7 cells (**Supplementary Figure 12**). To determine if similar changes in cellular metabolism occur in MCF-7 TaxR models, we performed Seahorse metabolic assays. The Cell Energy Phenotype test showed TaxR cells exhibit increased OCR and ECAR at baseline and stressed conditions relative to controls (**Figure 5C**), indicating increased mitochondrial respiration and glycolysis. To look at individual effects on OCR, we performed the Mito Stress test in MCF-7 TaxR models. Similar to cyto PELP1 expressing cells, TaxR cells showed significant increases in basal and maximal respiration compared to controls (**Figure 5D**). TaxR cells increased proton leak, spare respiratory capacity, and non-mitochondrial respiration, but not ATP production as observed in MCF-7 cyto PELP1 expressing cells. TaxR cells also displayed ~2-fold increase (p = 0.0006) in glucose uptake compared to controls (**Figure 5E**). Together, these data reveal that TamR and TaxR models phenocopy HIF-associated target gene expression and metabolic plasticity of MCF-7 cyto PELP1 expressing cells, and suggest PELP1 may be a key mediator in therapy resistance.

The PELP1/SRC-3 interaction was similarly increased in MCF-7 TaxR (**Figure 5F**, top) and TamR cells (**Figure 5F**, bottom). Additionally, CD44^hi^/CD24^lo^ ratios were increased in MCF-7 TaxR cells compared to parental controls (**Supplementary Figure 13**). To test the pharmacological effect of PFKFB3, PFKFB4, and SRC-3 inhibition, MCF-7 TaxR and TamR cells were seeded as tumorspheres and treated with PFK158, 5MPN, and SI-2. Both resistant models exhibited increased basal tumorsphere formation when compared to parental controls. 5MPN and SI-2 effectively decreased secondary tumorsphere formation by 71% and 75% in TaxR (**Figure 5G**), and 88% and 92% in TamR models (**Figure 5H**) compared to vehicle controls. PFK158 (PFKFB3 inhibitor) modestly decreased TaxR and TamR tumorspheres by 17% and 27%. These findings highlight the overlap of key players involved in PELP1-driven CSC biology and suggest that PFKFB4 and SRC-3 play a more significant role than PFKFB3 within resistant cell models.

We hypothesized that tam in combination with PELP1/SRC-3 complex inhibitors (i.e. SI-2 or 5MPN) would be more effective than either inhibitor alone. Combination treatments were evaluated in several cell lines. In MCF-7 PELP1 models, we tested tam/SI-2, tam/5MPN, and SI-2/5MPN combinations (**Figure 6A-6C**). Tam/SI-2 and tam/5MPN reduced tumorsphere formation in cyto PELP1 expressing cells by ~85% (p < 0.0001) and 80% (p < 0.0001) compared to vehicle. Single agent treatment with tam or SI-2 also reduced tumorspheres, but to a lesser degree than combinations. PFK158 co-treatment with tam was not more effective than tam alone and was not further pursued (**Supplementary Figure 14A**). Effective combinations were then tested in J110 cells (**Supplementary Figure 14B-14D**). Tam, SI-2, and 5MPN alone inhibited tumorspheres by 39, 41, and 28%, while co-treatment did not have dramatic effects. The SI-2/5MPN combination was most effective in J110 cells, and decreased tumorsphere formation by 60%, most likely because J110 cells are an SRC-3-derived transgenic mouse mammary tumor cell line (25).

**Figure 6.**
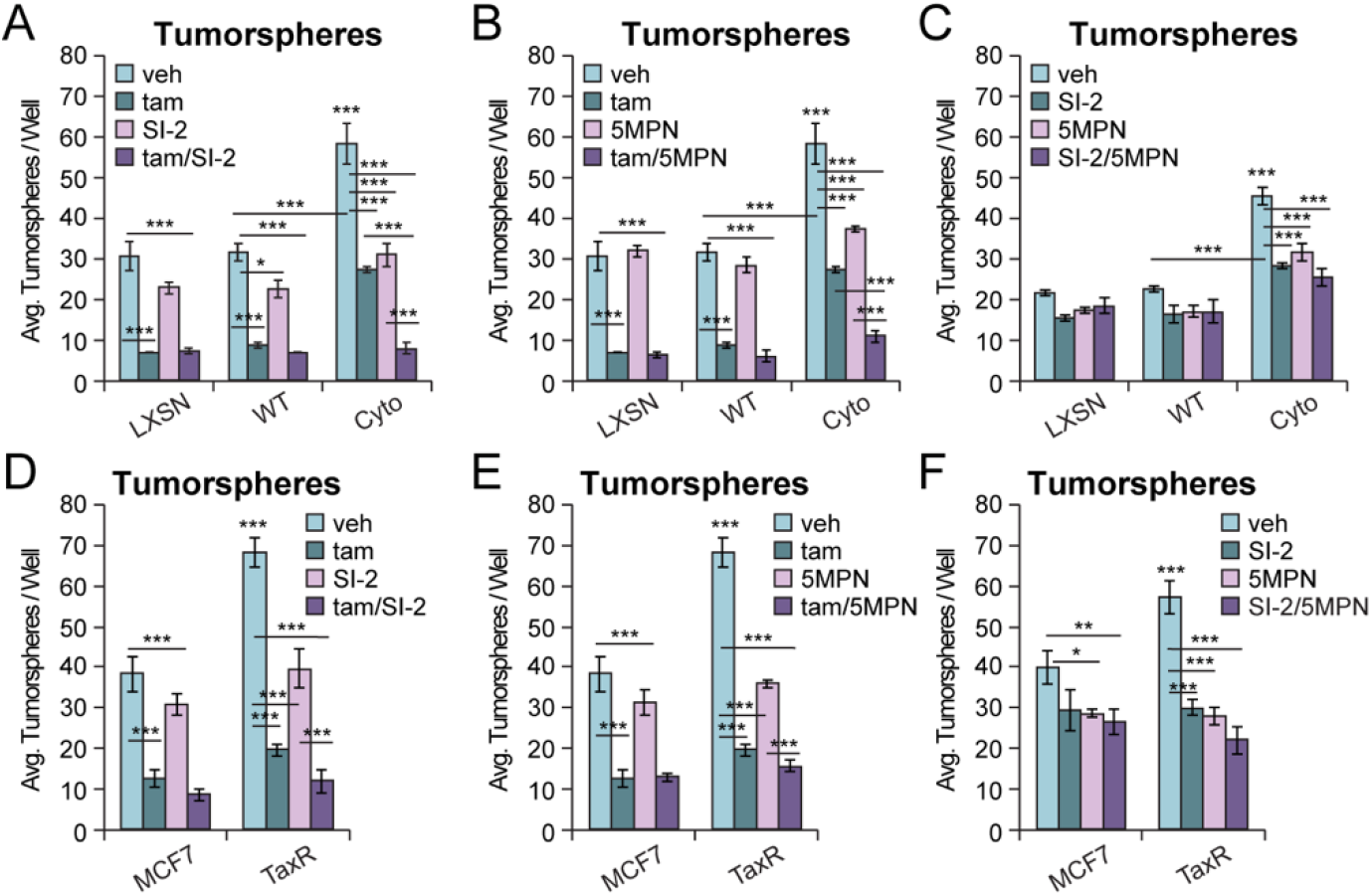
Endocrine therapies exhibit combinatorial effects with PELP1 complex inhibitors. Tumorsphere assays in MCF-7 PELP1 cells treated with: (**A**) tam/SI-2, (**B**) tam/5MPN, or (**C**) SI-2/5MPN. Tumorsphere assays in MCF-7 TaxR cells treated with: (**D**) tam/SI-2, (**E**) tam/5MPN, or (**F**) SI-2/5MPN. Concentrations: tam (100 nM), 5MPN (5 μM), SI-2 (100 nM). Graphed data represent the mean ± SD (n = 3). * p < 0.05, ** p < 0.01, *** p < 0.001.

Because PELP1 confers tamoxifen and Taxol resistance (**Figure 5A**), we also tested the effect of these agents in resistant cell models. Similar to observations in MCF-7 PELP1 models, tam co-treatments were more effective when combined with SI-2 or 5MPN in MCF-7 TaxR models (**Figure 6D, 6E**). The SI-2/5MPN combination was not more effective than individual agents in TaxR models (**Figure 6F**), suggesting that SRC-3 and PFKFB4 cooperation occurs in tam-sensitive models. Accordingly, SI-2/5MPN co-treatment in MCF-7 and T47D TamR models reduced tumorsphere formation by 77% (p < 0.0001) and 75% (p < 0.0001) (**Supplementary Figure 14E, 14F**).

To further explore the therapeutic potential of inhibitor combinations, we utilized pre-clinical patient-derived organoid models (PDxO;). Synergy screens were used to test combinations identified from **Figure 6** on proliferation using CellTiter Glo assays in PDxO models (HCI-003, −011, and −017). Zero Interaction Potency (ZIP) scores are shown in contour maps for tam/SI-2, tam/5MPN, and SI-2/5MPN treatments (**Figure 7A-7C, Supplementary Figure 15A-15C**), and indicate the percent a response is higher (>1) or lower (<1) than the expected response for the dose combination (δ-score). While the δ-score across the range of dose combinations tested were relatively weak, significant peaks of synergism (δ-score >5) were observed. The most synergistic area scores are summarized in **Figure 7D**. The tam/SI-2 δ-scores (~1 to 3) were the lowest and contour maps indicate antagonism. The SI-2/5MPN δ-scores (~5 to 13.5) are lower than the tam/5MPN scores (~12 to 27) suggesting the tam/5MPN combination is more effective at inhibiting PDxO proliferation. Next, we evaluated two of the ER+ PDxO models (HCI-003, HCI-017) for expression of PELP1, SRC-3, PFKFB3, PFKFB4, and ER mRNA and protein (**Supplementary Figure 15D, Figure 7E**). MCF-7 and T47D cell lines were included as controls. Both HCI-003 and HCI-017 express all proteins tested. Interestingly, PDxO models have higher levels of PFKFB proteins compared to MCF-7 and T47D cells. Next, we tested inhibitor combinations on CSC activity in PDxO models. PDxOs were grown to maturity, pre-treated for 3 days, then dissociated and seeded into tumorspheres in the presence of inhibitors. Individual treatments (tam, SI-2, 5MPN) reduced tumorsphere formation in both PDxO models by 36 to 62% (**Figure 7F-7H**). The tam/SI-2 combination was not more effective than individual treatment (**Figure 7F**). In contrast, tam/5MPN was more effective than tam or 5MPN alone and reduced tumorspheres by ~71% and ~90% in HCI-003 and HCI-017 (**Figure 7G**). SI-2/5MPN co-treatment was more effective than SI-2 or 5MPN alone and reduced tumorsphere formation by ~71% (p < 0.0001) and ~74% (p < 0.0001) in HCI-003 and HCI-017 (**Figure 7H**). These results demonstrate that blocking the PELP1/SRC-3 complex and associated binding partners is an effective approach to targeting CSC populations in multiple models of advanced breast cancer. Taken together, these studies provide promising alternative approaches to target non-ER mediators and overcome emergence of chemotherapy and endocrine resistance.

**Figure 7.**
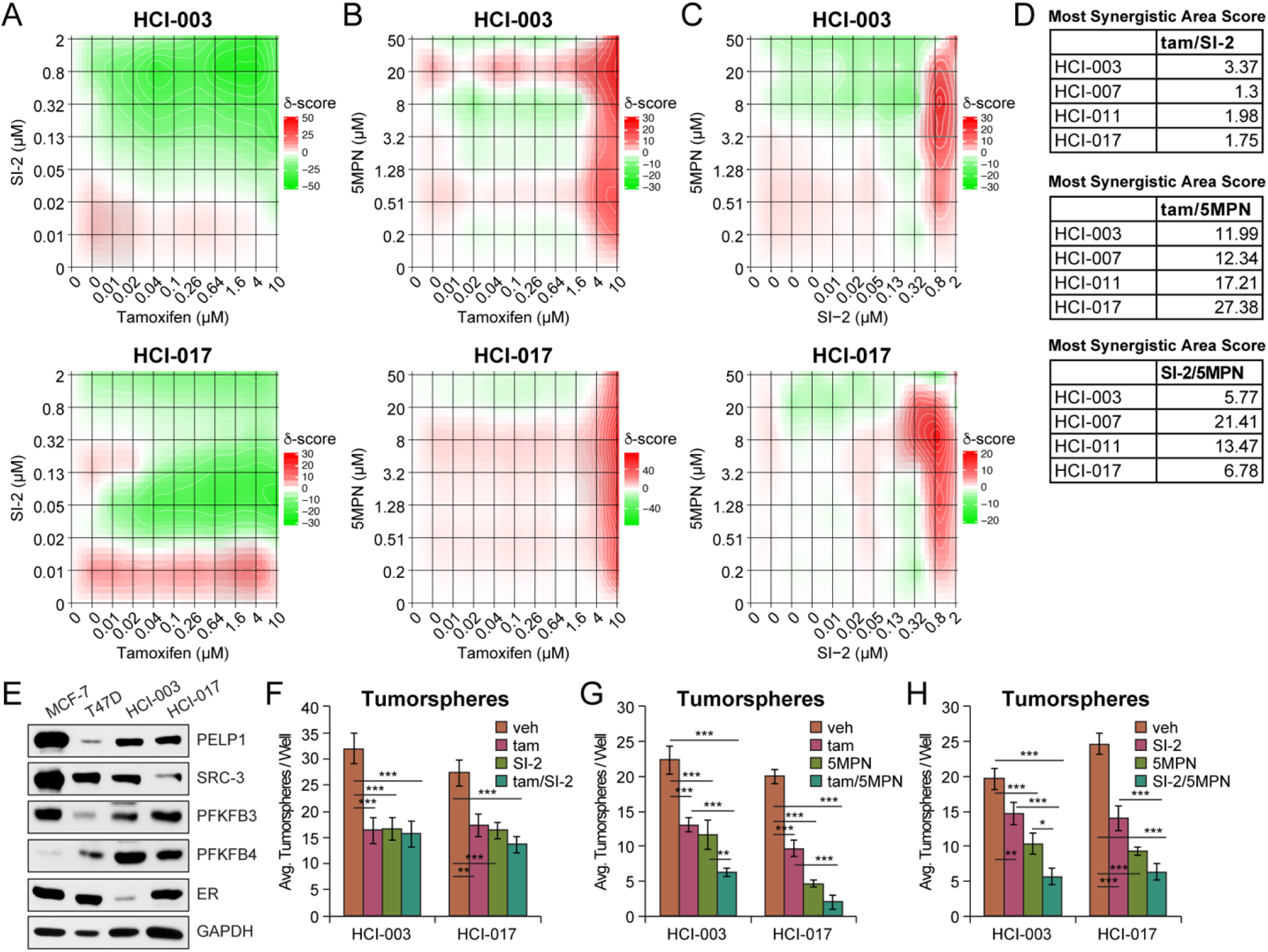
Co-treatments in preclinical ER+ PDxO models target CSCs. CellTiter Glo assays in HCI-003 and −017 PDxOs co-treated with **(A)** tam/SI-2, **(B)** tam/5MPN, or (**C**) SI-2/5MPN. (**D**) Tables summarizing most synergistic area scores from **7A-C and SFigure 15A-15C**. (**E**) Western blot of PELP1, SRC-3, PFKFB3, PFKFB4, and ER protein levels in HCI-003 and HCI-017. Tumorsphere assays in HCI-003 and HCI-017 PDxOs co-treated with (**F**) tam/SI-2, (**G**) tam/5MPN, or (**H**) SI-2/5MPN. Prior to assay, PDxO models were pre-treated with the indicated compounds for 3 days and subjected to continued treatment during the assay. Concentrations: tam (100 nM), 5MPN (5 μM), SI-2 (100 nM). Graphed data represent the mean ± SD (n = 3). * p < 0.05, ** p < 0.01, *** p < 0.001.

## DISCUSSION

The CSC hypothesis postulates that tumors contain a subset population (i.e. CSCs) that share properties of normal stem cells including self-renewal, differentiation, and capacity to repopulate the heterogeneous tumor (9). CSCs are proposed to have heightened resistance to cancer therapies due to their relative quiescent state (26), enabling this population to evade standard of care treatments that target proliferating bulk tumor cells. Herein, we sought to define mechanisms of SR co-activator driven CSC survival and expansion in ER+ breast cancer. We conclude that SR co-activator complexes enhance CSC activity and therapy resistance by promoting metabolic plasticity. Inhibiting these complexes and/or associated binding partners in combination with endocrine therapies may be an effective strategy to block CSC survival and self-renewal, and breast cancer progression.

Our findings further implicate PELP1/SRC-3 complexes as mediators of CSC activity. We observed similarities in gene expression, cell metabolism, and sensitivity to inhibitors of PELP1 binding partners in endocrine and chemotherapy resistant ER+ cell lines. Although PELP1 expression contributes to cell survival in response to Taxol in TNBC (14), our studies are the first to demonstrate enhanced Taxol tolerance in the context of cyto PELP1 in ER+ breast cancer. Our results in TaxR models highlight the impact of targeting PELP1 binding partners involved in PELP1-mediated CSC self-renewal (**Figure 5**). Mesenchymal stem cells (27) and ovarian cancer cells (28) achieve Taxol resistance by shifting to G0 and entering quiescence. PELP1 is a substrate of CDKs and modulates G1/S cell cycle progression (29). PELP1 may confer Taxol resistance in part through cell cycle regulation, albeit further studies are needed to define cytoplasmic PELP1-specific contributions in this context.

Contributing factors to CSC survival include metabolic plasticity, which enables adaptation to diverse tumor environments. For example, inhibition of glycolysis reduces breast and lung CSCs (30). Glycolytic reprogramming has been documented in breast cancer cells during EMT, resulting in acquisition of CSC-like characteristics and tumorigenicity (31). In contrast, breast CSCs utilize oxidative phosphorylation (OXPHOS) as their primary metabolic program (32). Bulk tumor cells depend chiefly on glycolysis, whereas tumors enriched for breast CSCs rely mainly on OXPHOS (33). RNA-seq analysis indicated cytoplasmic PELP1 imparts increased HIF-activated pathways under normoxic 3D conditions to enrich for CSCs. ChIP assays demonstrated *EPAS1* (i.e. HIF-2α) recruitment to HRE regions of the PELP1 promoter in TNBC cells (34). Thus, PELP1-induced HIF pathways may serve as a feed-forward mechanism to drive metabolic genes programs. PFKFB3 and PFKFB4 are required for glycolytic response to hypoxia via HIF-1α activation (20). We demonstrated that cyto PELP1 expressing cells increased glycolysis and mitochondrial respiration. Additional studies are needed to define the bioenergetics driving this plasticity. PFKFB4-mediated SRC-3 Ser857 phosphorylation has essential functions in lung and breast cancer metastasis and metabolism (35). Phosphorylation of SRC-3 Ser857 promotes SRC-3 association with transcription factor *ATF4* to mediate non-oxidative pentose phosphate pathway and purine synthesis. This study (35) did not evaluate SRC-3 in the context of CSCs, although SRC-3 has been linked to CSC activity (10,11). Our IPA studies also identified ATF4 pathway activation (**Figure 1**); upregulation of *ATF4* could explain the correlation between PFKFB4 and PELP1/SRC-3-driven CSCs.

PFKFB inhibitors are emerging as promising treatments in endocrine and chemotherapy-resistant ER+ breast cancer (36). PFKFB3 inhibitor, PFK158, displays broad anti-tumor and immunomodulatory effects in human and preclinical mouse models (37) and was evaluated in a Phase I clinical trial with no significant adverse effects (38). The prognostic value of PFKFB4 expression was evaluated in 200 tumor samples from stage I to III breast cancer patients. Similar to our METABRIC analysis (**Supplementary Figure 10**), elevated PFKFB4 expression was associated with poor disease-free survival and overall survival in ER+, HER2+, or TNBC patients (39). PFKFB4 inhibitors (e.g. 5MPN) have not yet moved to clinical trials. Studies have suggested correlative and mechanistic links between PFKFBs and CSCs. *PFKFB3* was upregulated in a CD44^hi^CD24^lo^ gene signature correlated to risk of distant metastasis and poor outcome in breast cancer patients (40). A cleaved product of CD44 (CD44ICD) promoted breast cancer stemness via PFKFB4-mediated glycolysis (41). Notably, 5MPN treatment suppressed CD44ICD-induced tumorigenesis. We have further implicated PFKFBs as drivers of CSC activity by demonstrating 5MPN reduces tumorspheres as a single agent or in combination treatments in multiple ER+ breast cancer models, including treatment resistant cells (TaxR, TamR), murine tumor cells, and pre-clinical PDxOs. Our data shows that treatment with 5MPN in combination with SI-2 or tam inhibits PDxO proliferation (**Figure 7**), but importantly also targets the CSC population, Studies in breast cancer patients indicate that EMT and CSC markers are present in CTC populations, which have high metastatic potential (42). Our MIND xenografts demonstrate PFKFB4 knockdown does not have an effect on primary tumor burden, but reduces CTC populations (**Figure 4**). These data suggest PFKFB4 inhibition is an effective strategy for targeting CSCs and CTCs in ER+ breast cancer. Future work should involve assessing overlap between PFKFB4-modulated CSC and CTC populations by evaluating the impact of 5MPN inhibitor combinations *in vivo*.

To evaluate the impact of SR co-activators on CSCs, it will be important that future studies consider SR-driven contributions. Breast CSCs are reported to be mostly ER negative (43), which may explain their poor response to anti-estrogens. However, SR+ cells contribute to CSC biology through SR-dependent (namely PR) paracrine factors (44). For example, breast CSC self-renewal was stimulated after anti-estrogen treatment of breast cancer cells or PDX models (45,46). These studies suggest anti-estrogen therapies may initially slow tumor growth, but concurrently evoke plasticity and CSC activity in non-proliferative tumor cells. Notably, PELP1-containing complexes include ER and PR (47). PRs but not ER are potent drivers of stem and progenitor cell expansion in normal and neoplastic breast tissues (48). We have recently defined a requirement for phosphorylated and inducible PR in CSC biology (49), insulin hypersensitivity, and tam resistance in ER+ breast cancer (13). CSC outgrowth in therapy resistant ER+/PR-low breast cancer models is blocked by PR knockdown or antiprogestins (13). These findings suggest PELP1/SRC-3 complexes enable constitutive SR activity in sub-populations that easily bypass endocrine therapies. Antagonizing estrogen signaling may select for cells that display ligand-independent ER and PR, resulting in increased proportions of breast CSCs, and subsequently promote metastasis. Therefore, treatment should include endocrine therapy in combination with targeted therapies that block mediators of CSC survival and self-renewal as defined herein (i.e. PFKFBs).

## CONCLUSION

Our work demonstrates that targeting SR co-activators and associated binding partners involved in driving CSC survival, self-renewal, and metabolic plasticity may impede breast cancer progression and has the potential to lead to improved outcomes. Identifying the mechanisms that mediate recurrent ER+ tumor cell populations (e.g. CSCs, CTCs) will enable specific targeting within heterogeneous breast tumors to overcome endocrine and chemotherapy resistance.

## ACKNOWLEDGEMENTS

We thank Bruce Lindgren for biostatistics support, and the Masonic Cancer Center Biostatistics and Bioinformatics, Analytical Biochemistry, and Flow Cytometry shared resources. We also thank Zohar Sachs and Michael Franklin for critical reading of this manuscript.

