## Supplementary FIgures for "PELP1/SRC-3-dependent regulation of metabolic kinases drives therapy resistant ER+ breast cancer"

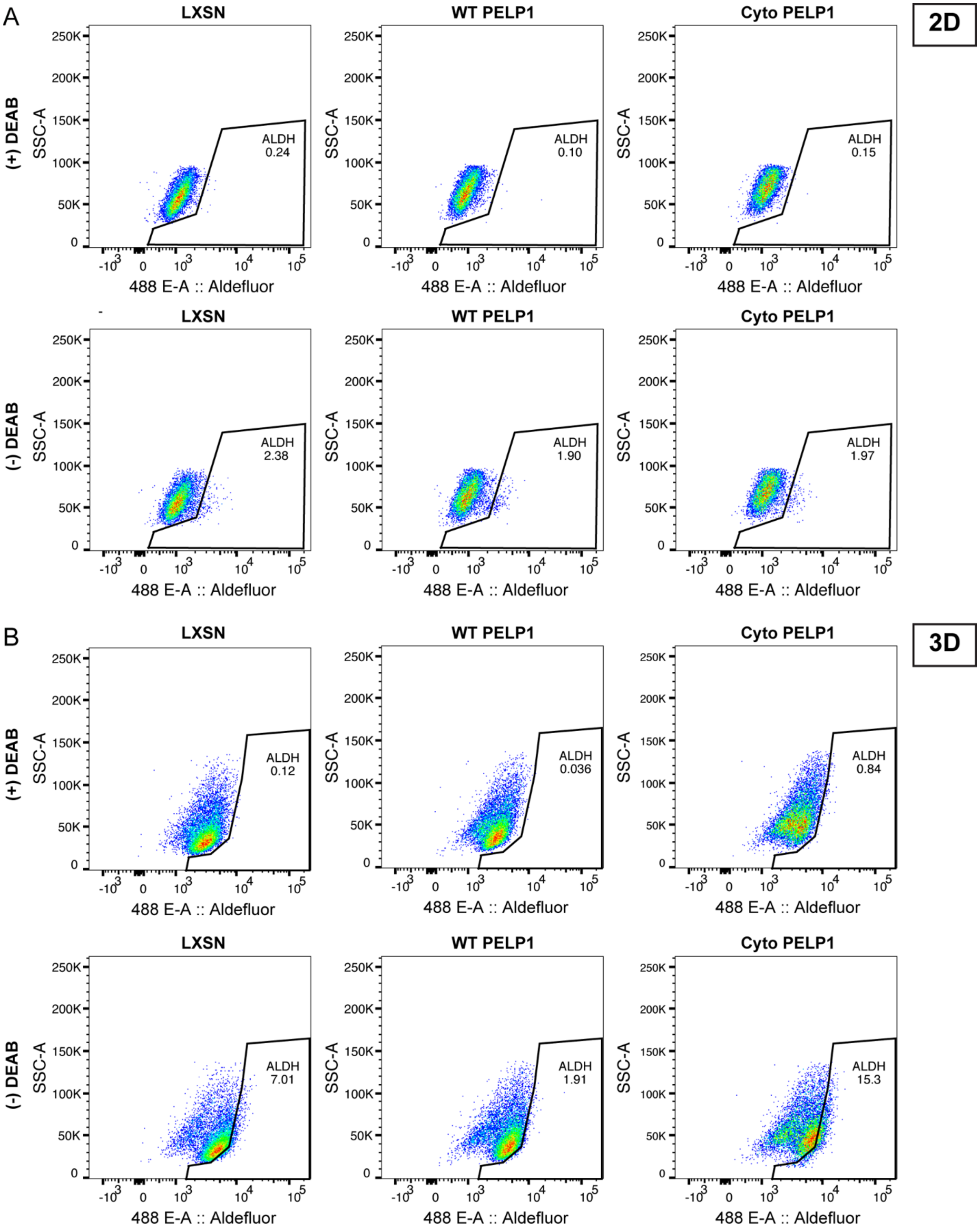


**Supplementary Figure 1.** Representative flow cytometry dot plots shown for ALDH activity in MCF-7 PELP1 cells cultured in (**A**) 2D (adherent) or (**B**) 3D (tumorsphere) conditions.

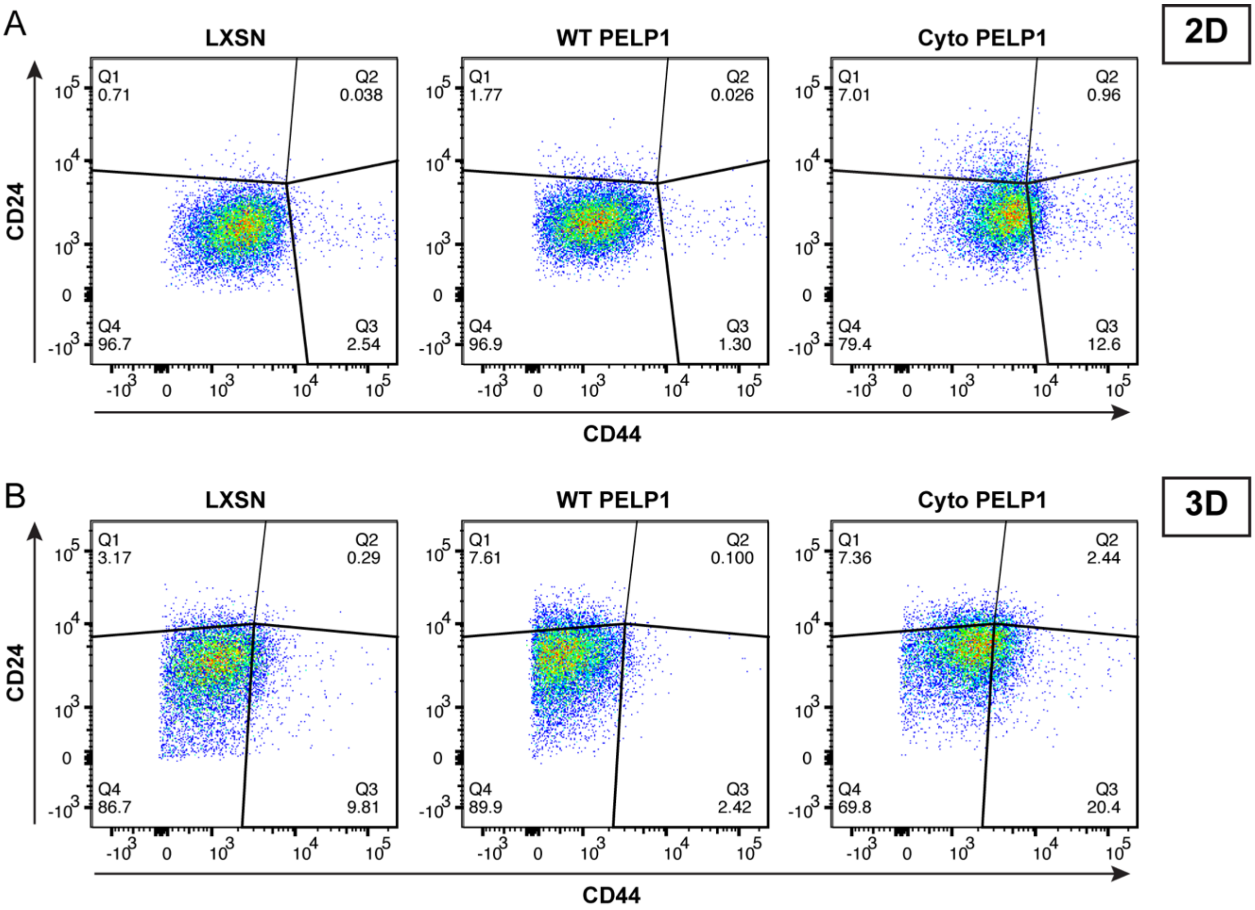


**Supplementary Figure 2.** Representative flow cytometry dot plots shown for CD44/CD24 populations in MCF-7 PELP1 cells cultured in (**A**) 2D (adherent) or (**B**) 3D (tumorsphere) conditions.

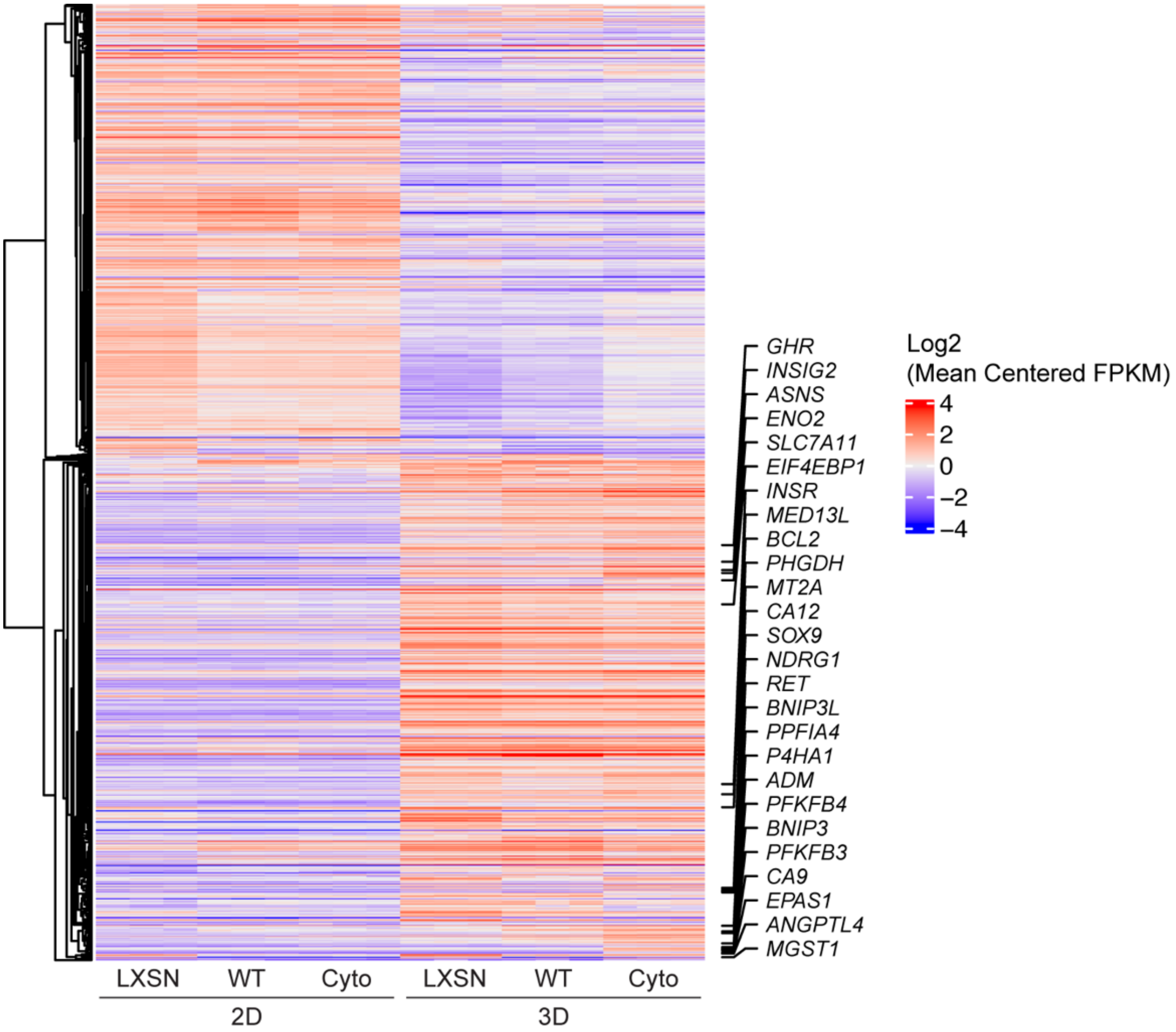


**Supplementary Figure 3.** Supervised heat map of differentially expressed genes in MCF-7 PELP1 cells (LXSN, WT PELP1, cyto PELP1) in 3D vs. 2D culture. Genes were hierarchically clustered using average clustering method and Pearson correlation as distance in R ComplexHeatmap package (PMID: 27207943). Columns are grouped by biological replicates for display purposes. PELP1 signature genes are marked.

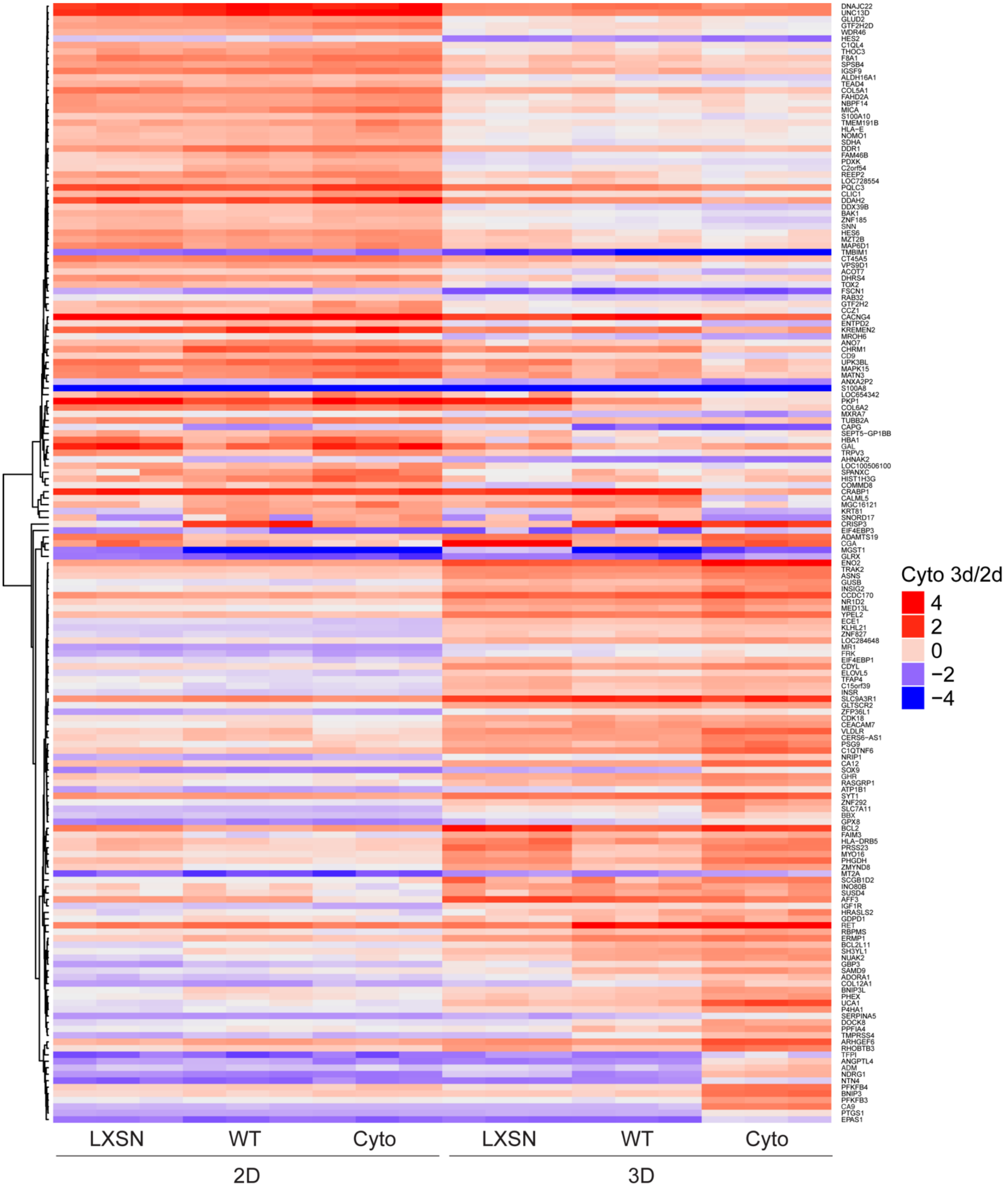


**Supplementary Figure 4.** Supervised heat map of differentially expressed genes in MCF-7 cyto PELP1 (3D vs. 2D culture).

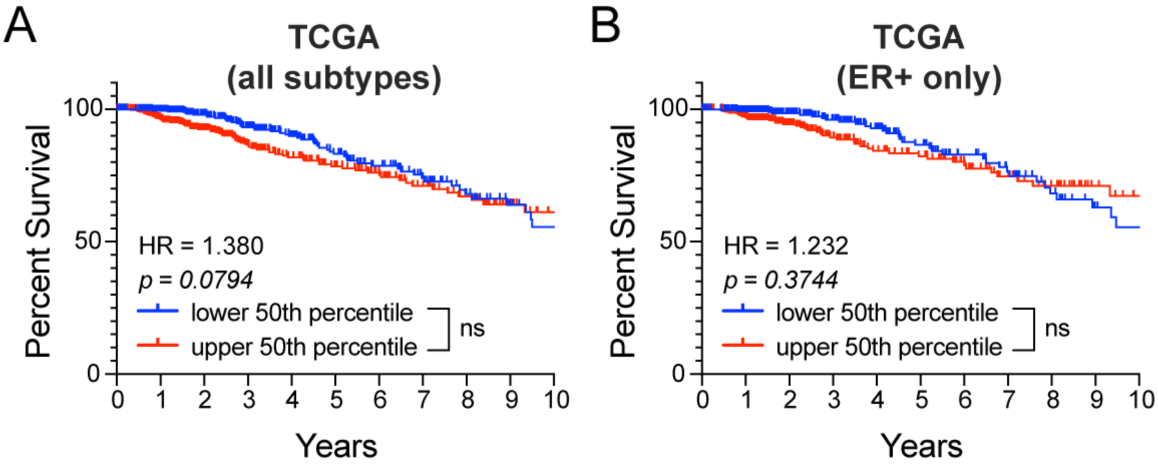


**Supplementary Figure 5.** Kaplan-Meier curves shown for upper and lower 50$^{th}$ percentile of the cyto PELP1 gene signature in the TCGA (**A**) all subtypes and (**B**) ER+ only patient cohorts.

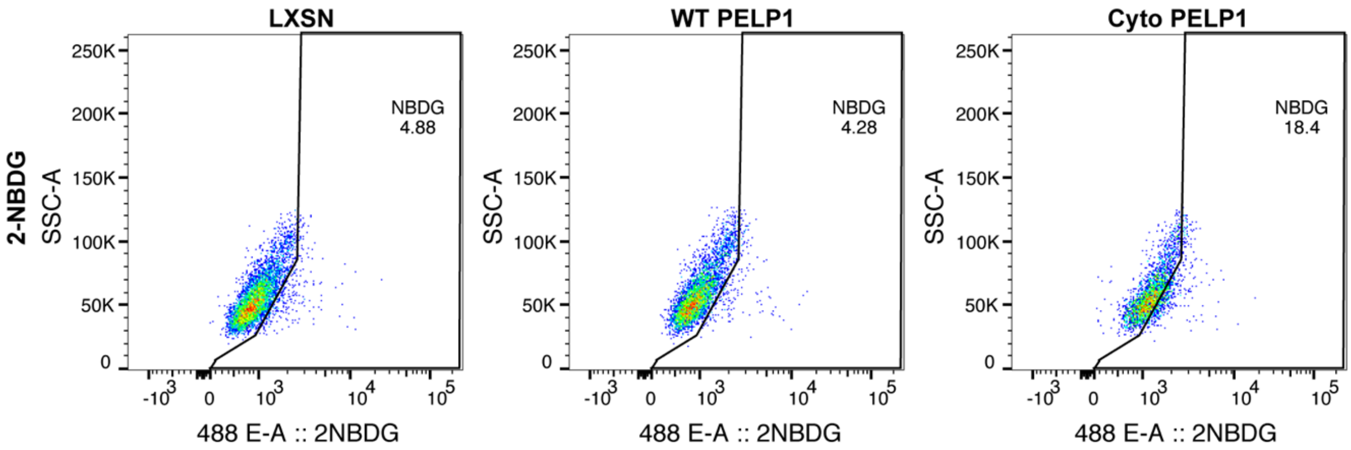


**Supplementary Figure 6.** Representative flow cytometry dot plots shown for 2-NBDG glucose uptake assays in MCF-7 PELP1 cells.

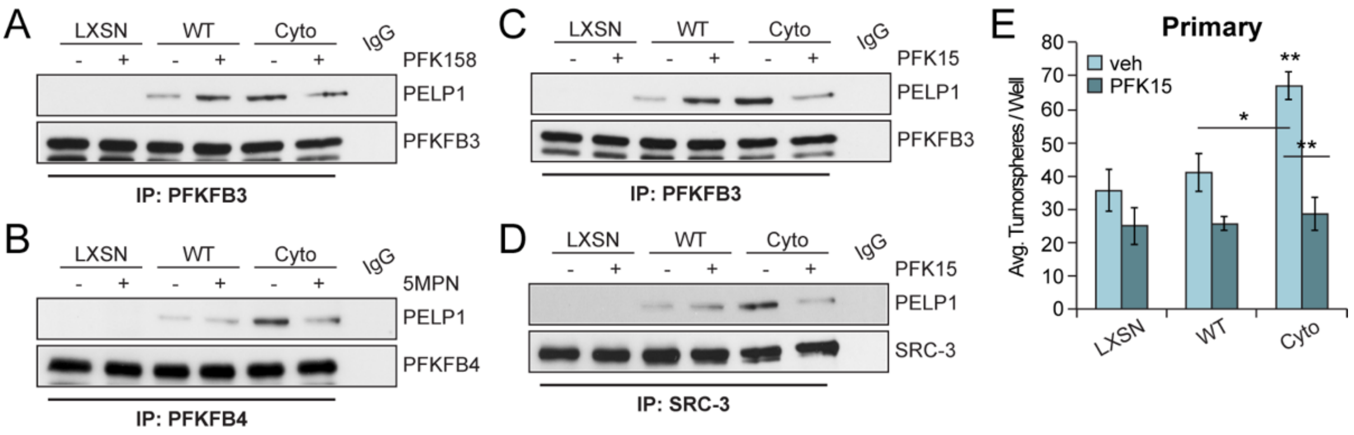


**Supplementary Figure 7.** Co-immunoprecipitation of (**A**) PELP1 and PFKFB3 treated with vehicle (DMSO) or PFK158 (100 nM) and (**B**) PELP1 and PFKFB4 treated with vehicle or 5MPN (5 μM) in MCF-7 PELP1 cells. Co-immunoprecipitation of (**C**) PELP1/PFKFB3 and (**D**) PELP1/SRC-3 in MCF-7 PELP1 cells treated with vehicle or PFK15 (100 nM). (**E**) Primary tumorsphere assays in MCF-7 PELP1 cells treated with vehicle or PFK15. Graphed data represent the mean ± SD (n = 3). * $p < 0.05$, ** $p < 0.01$, *** $p < 0.001$.

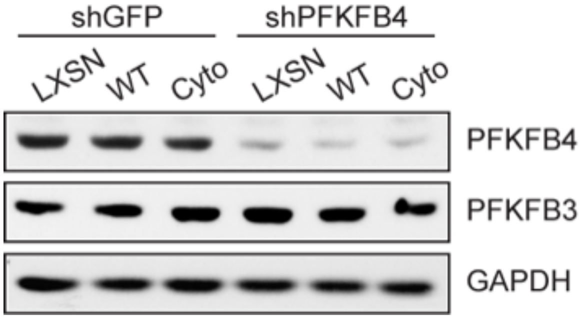


**Supplementary Figure 8.** Western blot showing PFKFB4 knockdown in MCF-7 PELP1 models.

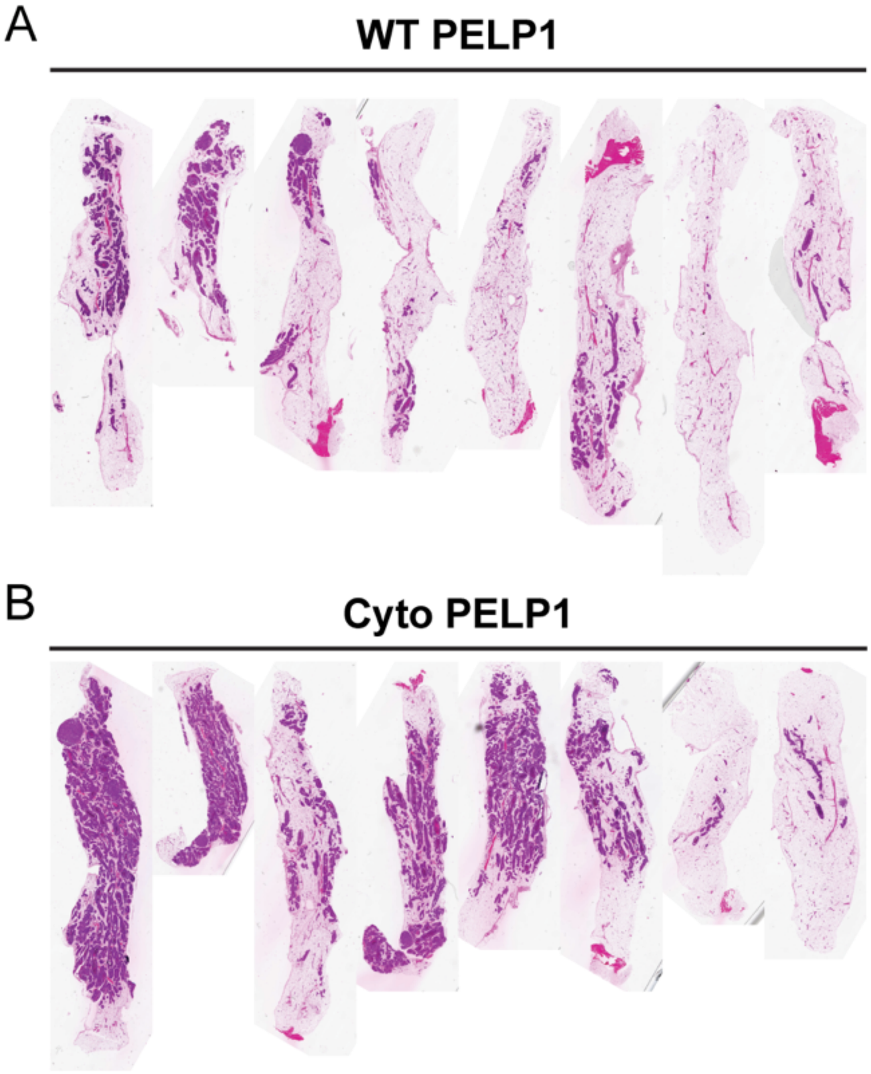


**Supplementary Figure 9.** H&E stains from MIND glands in (**A**) WT PELP1 and (**B**) cyto PELP1.

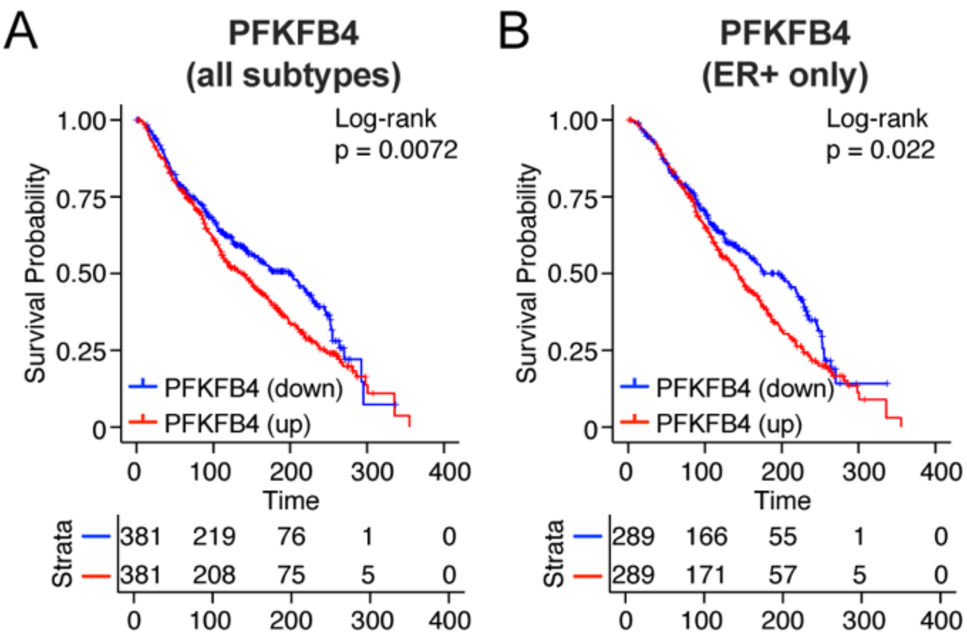


**Supplementary Figure 10.** Kaplan-Meier curves shown for PFKFB4 expression in the METABRIC (**A**) all subtypes and (**B**) ER+ only patient cohorts.

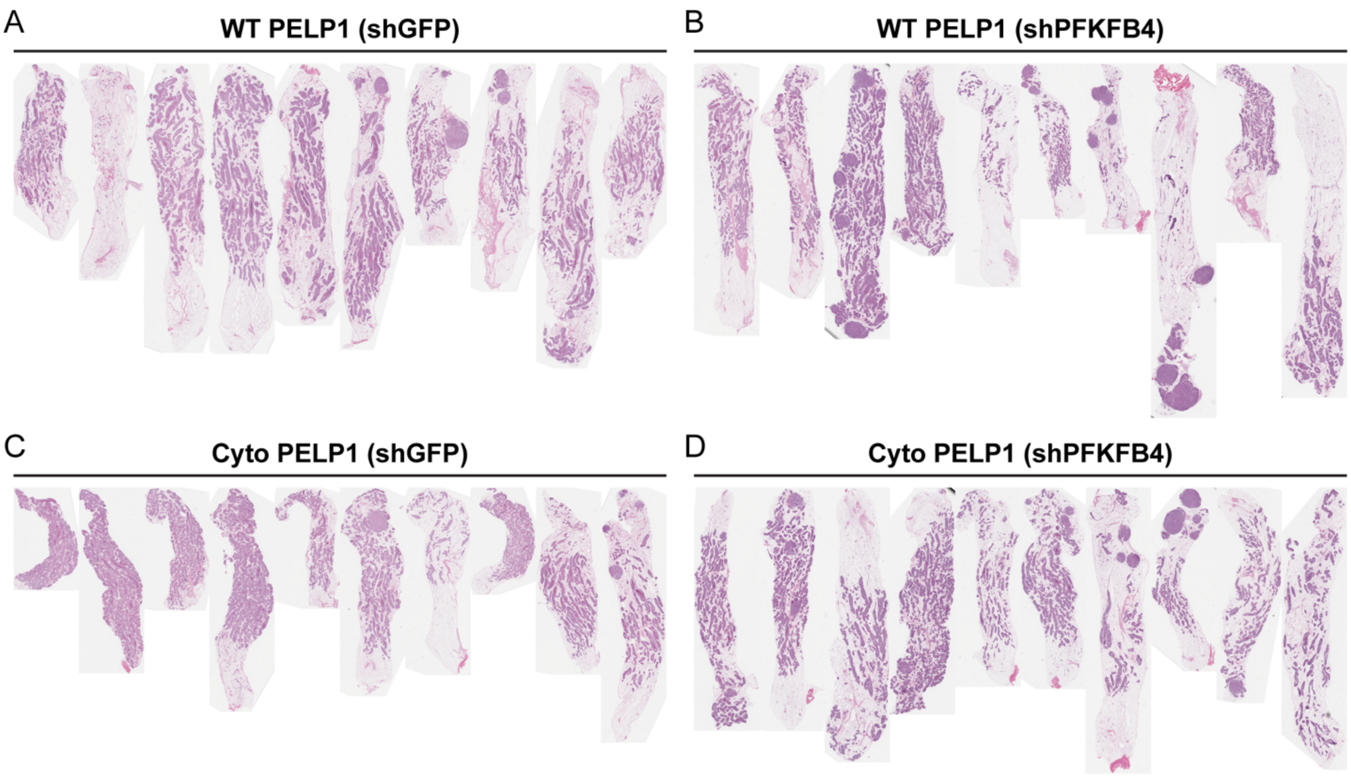


**Supplementary Figure 11.** H&E stains from MIND glands in (**A, B**) WT PELP1 (shGFP, shPFKFB4) and (**C, D**) cyto PELP1 (shGFP or shPFKFB4).

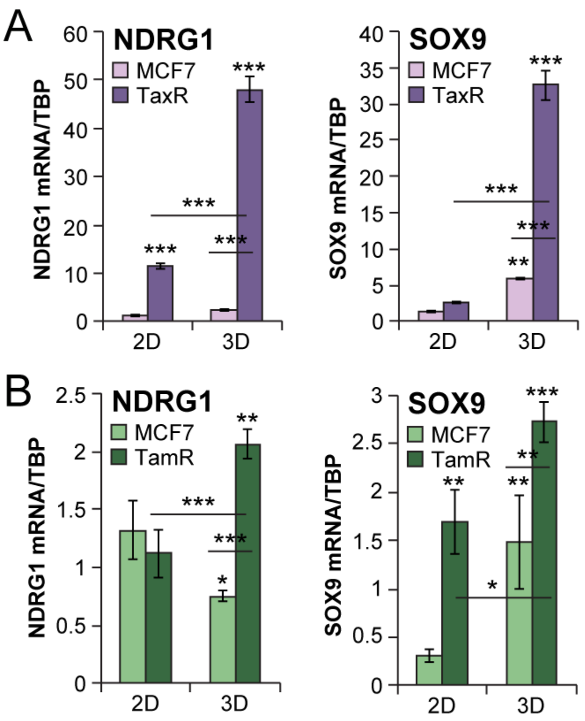


**Supplementary Figure 12.** RNA-seq gene validation. mRNA levels of HIF-activated metabolic genes (*NDRG1*) and HIF-activated stem cell genes (*SOX9*) in (**A**) MCF-7 TaxR and (**B**) MCF-7 TamR cells.

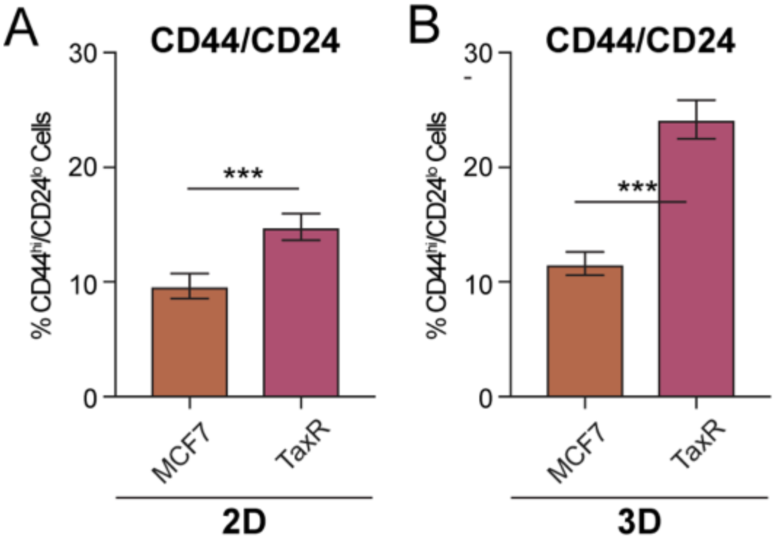


**Supplementary Figure 13.** $CD44^{hi}/CD24^{lo}$ populations in (**A**) 2D (adherent) or (**B**) 3D (tumorsphere) conditions in MCF-7 TaxR cells. Graphed data represent the mean ± SD (n = 3). * $p < 0.05$, ** $p < 0.01$, *** $p < 0.001$.

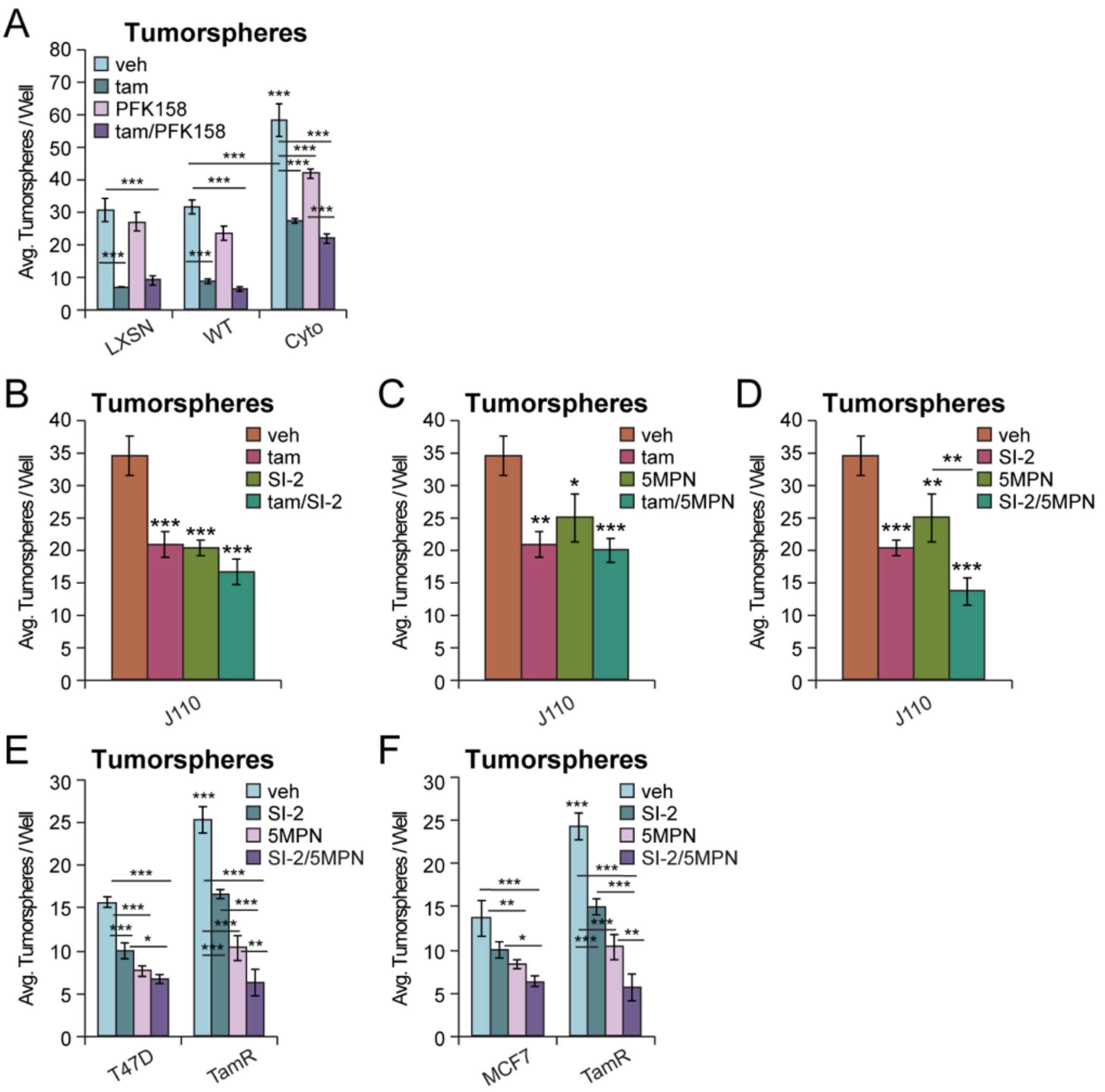


**Supplementary Figure 14.** Tumorsphere assays in MCF-7 PELP1 cells treated with (**A**) tam/PFK158. Tumorsphere assays in J110 cells treated with: (**B**) tam/SI-2, (**C**) tam/5MPN, (**D**) SI-2/5MPN. Tumorsphere assays in (**E**) T47D TamR and (**F**) MCF-7 TamR cells treated with SI-2/5MPN. Concentrations used: tam (100 nM), PFK158 (100 nM), 5MPN (5 µM), SI-2 (100 nM). Graphed data represent the mean ± SD (n = 3). * $p < 0.05$, ** $p < 0.01$, *** $p < 0.001$.

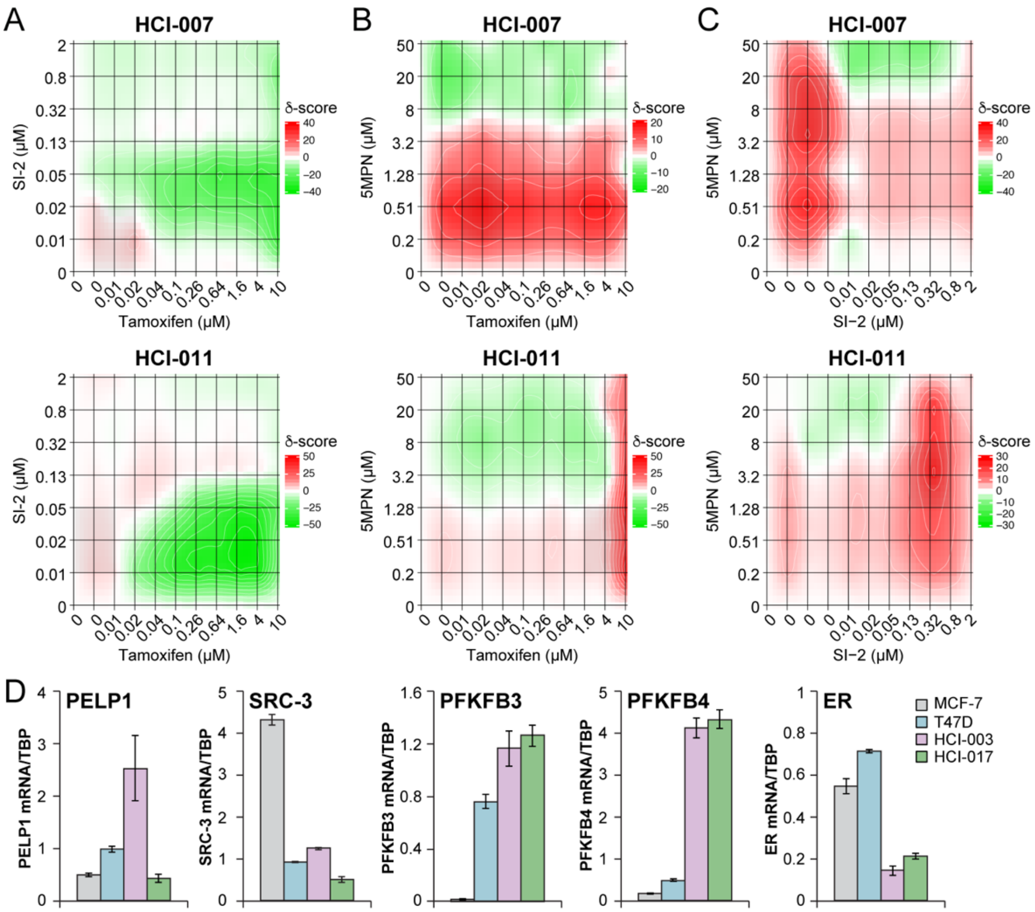


**Supplementary Figure 15.** CellTiter Glo assays in HCI-007 and -011 PDxOs co-treated with (**A**) tam/SI-2, (**B**) tam/5MPN, or (**C**) SI-2/5MPN. (**D**) PDxO characterization of PELP1, SRC-3, PFKFB3, PFKFB4, and ER mRNA levels in HCI-003 and HCI-017.
