## Supplementary Methods for "PELP1/SRC-3-dependent regulation of metabolic kinases drives therapy resistant ER+ breast cancer"

**General Reagents.** Hydrocortisone (Sigma) and tamoxifen (tam; Sigma) were prepared in ethanol (EtOH). Epidermal growth factor (EGF; Sigma) was prepared in 0.1% BSA. Paclitaxel (Taxol; Thermo Fisher), PFK15 (Tocris), PFK158 (Cayman Chemicals), 5MPN (12), and SI-2 (Tocris) were prepared in DMSO.

**Stable Cell Line Generation.** Stable shPFKFB4 (clones TRCN0000037764, -37767) cells were created by transducing MCF-7 PELP1 models with pLKO.1 lentivirus and maintained as described above with 0.5 µg/ml puromycin (MP Biomedicals).

**Tumorsphere Assays.** Single cell suspension was filtered through a 40-µm sieve (BD Falcon) and seeded in ULA plates. Cells were grown in supplemented MEBM media. For secondary tumorspheres, primary tumorspheres were dissociated in 0.25% trypsin-EDTA. Cells were plated as described above in supplemented MEBM media. Tumorspheres were allowed to form for 7-10 days. Tumorspheres were analyzed by total number and scored by manual counting using a scaled grid. Data are presented as the average ± SD of three independent measurements.

**Flow Cytometry.** An ALDEFLUOR assay kit (Stem Cell Technologies) was used to measure aldehyde dehydrogenase (ALDH) activity. For 2D conditions, cells were plated into growth media. For 3D conditions, cells were plated into ULA plates in supplemented MEBM media. Cells were dissociated using Accutase (Life Technologies) and resuspended ($5 \times 10^5$ cells) in ALDEFLUOR buffer. ALDEFLUOR reagent was added, and then half of the cells were transferred to a control tube containing DEAB. Cells were incubated at 37 °C for 45 min, washed, and subjected to flow cytometry (BD LSRII H4760). Sorting gates were established using DEAB-treated cells.

For CD44/CD24 detection, cells were dissociated with Accutase. Cells ($5 \times 10^5$) were resuspended in FACS buffer (D-PBS containing 2% FBS) with APC-CD44 (1:20; BD Pharmingen) and CD24-PE (1:50; BD Pharmingen) conjugated antibodies and incubated at 4 °C for 30 min. Cells were washed, resuspended in cold FACS buffer, and subjected to flow cytometry. Data were plotted as CD24-PE versus CD44-APC to identify populations based on single stained controls.

**Glucose Uptake.** Glucose uptake was measured using 2-NBDG (Thermo Fisher). Cells were incubated in glucose-free DMEM containing 1% HEPES (Gibco) for 15 min and then treated with

2-NBDG for 25 min at 37 °C. Cells were dissociated using trypsin, washed, and resuspended in cold FACS buffer (D-PBS containing 2% FBS). 2-NBDG fluorescence was quantified by flow cytometry, and sorting gates were established using untreated control cells.

**Seahorse Assays.** Seahorse XFe96 Analyzer (Agilent) was used to measure ECAR and OCR levels. MCF-7 PELP1 cells were seeded into Agilent XF96 culture plates (2.5 x $10^4$ cells/well with 22.4 μg/ml Cell Tak) and incubated at 37 °C for 24 h. Cells were washed 2x and incubated in Seahorse XF DMEM media (2 mM L-glutamine, 11.11 mM glucose, 1.0 mM sodium pyruvate; pH 7.4) at 37 °C in a $CO_2$-free incubator for 45 min. ECAR and OCR were detected under basal conditions followed by addition of 0.5 μM FCCP and 2 μM oligomycin using the XF Cell Energy Phenotype and Mito Stress Test (Agilent). Protein concentrations were measured by BCA Assay following addition of RIPA-lite (25 μl/well). ECAR and OCR data were normalized to total protein concentration per well. Data are presented as the average ± SD of experimental triplicates. MCF-7 TaxR cell conditions: 1 x $10^4$ cells/well, Seahorse XF DMEM media (2 mM L-glutamine, 5.55 mM glucose, 1 mM sodium pyruvate; pH 7.4), FCCP (0.25-0.5 μM), and oligomycin (2 μM).

**Real-Time Quantitative-PCR (RT-qPCR).** RNA was extracted using TriPure Isolation Reagent (Roche) and isopropanol precipitation. RNA (1000 ng) was reverse transcribed to cDNA using qScript cDNA SuperMix (Quanta BioSciences). qPCR was performed using Light Cycler FastStart DNA Master SYBR Green I (Roche) on a Light Cycler 96 Real-Time PCR System (Roche). Conditions: initial denaturation at 95°C (10 min), denature at 95°C (10 sec), anneal at 60°C (10 sec), and extension at 72°C (5 sec) for 45 cycles. Gene levels were normalized to housekeeper genes and represent the average ± SD of three independent measurements

**RNA-Sequencing.** For 3D conditions, cells were plated as described above. 50 base pair paired-end sequencing was performed using Illumina HiSeq 2500 system. On average >12 million reads were sequenced per sample. Each sample was aligned using the Tophat aligner (v 2.0.13). Samtools (v 1.0_BCFTools_HTSLib) was used to sort and index bam files. Cuffquant (Cufflinks v 2.2.1) was used to generate transcript abundance files. Once samples were mapped and abundance estimate files were completed, Cuffnorm (Cufflinks v 2.2.1) was used to generate a table of Fragments Per Kilobase Of Exon Per Million Fragments Mapped (FPKM) values for genes within each sample. For each gene, FPKM values were log transformed and mean centered. Genes that varied <0.1 standard deviation within the sample set were removed from further analysis. A simple t-test with Benjamini-Hochberg correction where genes with q-values <0.05

and minimum of 2-fold expression change were used to identify differentially expressed genes between conditions.

**Cell Lysate Preparation.** Cells were harvested in RIPA-lite lysis buffer [150 mM NaCl, 6 mM $Na_2HPO_4$, 4 mM $NaH_2PO_4$, 2 mM EDTA, 100 mM NaF, 1% Triton-X 100, 1X complete mini protease inhibitors (Roche), 1X PhosSTOP (Roche), and supplemented with 1 mM PMSF, 5 mM NaF, 0.05 mM $Na_3VO_4$, 25 mM beta glycerophosphate (BGP), and 20 µg/ml aprotinin].

**Co-Immunoprecipitation Assays.** Cells were harvested in ELB lysis buffer [50 mM HEPES, 0.1% NP-40, 250 mM NaCl, 5 mM EDTA, 1X complete protease inhibitors (Roche), 1X PhosSTOP (Roche), and supplemented with 1 mM PMSF, 1mM NaF, 0.5 mM $Na_3PO_4$, 25 mM BGP, and 20 µg/ml aprotinin]. 1000 µg lysate (1 mg/ml) was incubated with 1 µg of the indicated antibody overnight at 4°C. Immunocomplexes were isolated with protein G agarose (Roche) for 2 h at 4°C. Resin was collected and washed with cold ELB buffer. Immunocomplexes were eluted with sample buffer, resolved by SDS-PAGE, and analyzed by Western blot.

**Immunoblotting.** Antibodies used: SRC-3 (5E11, Cell Signaling), GAPDH (0411, Santa Cruz Biotechnology), PELP1 (A300-180A, Bethyl Labs), PFKFB3 (D7H4Q, Cell Signaling), PFKFB4 (PA5-28648, Thermo Fisher), ERα (F-10, Santa Cruz Biotechnology), goat anti-rabbit IgG-HRP (BioRad), and goat anti-mouse IgG-HRP (BioRad).

**Image Analysis.** Each H&E stained section was analyzed using QuPath 0.2.1. Two pixel classifiers were used to identify tissue and tumor regions, respectively. Tumor regions were split into individual areas and measured, excluding areas below 312.5 $um^2$ and filling holes up to 62.5 $um^2$. The total area of the resulting tumor regions was divided by the area of the tissue section times 100 to obtain the percentage coverage. Code and QuPath classifiers are available at https://github.com/tp81/ostrander-2020.

**CTC Soft Agar Assays.** Fresh mouse blood samples were processed using Isolymph/Ficoll-Paque (Sigma). Buffy coat containing circulating tumor cells was seeded into DMEM containing 5% FBS, 1X sterile low melt agarose (Thermo Fisher), and 1X penicillin streptomycin. Colonies were grown for 14 days at 37 °C. Data are presented as the average ± SD of five independent measurements.

**Public Data Mining.** METABRIC and TCGA datasets were downloaded from the cBioPortal website for genes indicated in each signature. R Studio (v 1.1.383) was utilized for data processing. Values obtained for each dataset were log2 transformed, averaged across each patient, and nominated as the average signature score. The median for the average signature score was calculated for all patients. Patients with an average signature score higher than the median were nominated as upper 50$^{th}$ percentile; average signature score lower than the median were nominated as lower 50$^{th}$ percentile. Kaplan-Meier curves were plotted using PRISM (v 8.3.1).
