## Supplementary Tables for "PELP1/SRC-3-dependent regulation of metabolic kinases drives therapy resistant ER+ breast cancer"

**Supplementary Table 1:** IPA pathways (all genes)

**Supplementary Table 1**

| Upstream Regulator | Predicted Activation State | Activation z-score | p-value of overlap |
|---|---|---|---|
| beta-estradiol | Activated | 4.076 | 2.12E-30 |
| IFNG | Activated | 4.106 | 1.66E-26 |
| TNF | Activated | 3.149 | 3.74E-20 |
| IL1B | Activated | 3.759 | 4.78E-19 |
| EGF | Activated | 2.804 | 1.27E-16 |
| lipopolysaccharide | Activated | 4.822 | 1.41E-15 |
| FGF2 | Activated | 2.796 | 1.2E-14 |
| tretinoin | Activated | 3.726 | 1.42E-14 |
| IL6 | Activated | 2.789 | 3.63E-14 |
| FSH | Activated | 2.004 | 1.99E-13 |
| IL2 | Activated | 2.303 | 3.96E-13 |
| JUN | Activated | 2.105 | 4.07E-13 |
| STAT3 | Activated | 2.292 | 1.29E-12 |
| HGF | Activated | 2.742 | 1.71E-12 |
| cigarette smoke | Activated | 2.339 | 2.99E-12 |
| NFkB (complex) | Activated | 4.301 | 1.22E-11 |
| PDGF BB | Activated | 3.512 | 1.43E-11 |
| CTR9 | Activated | 2.333 | 1.56E-11 |
| IGF1 | Activated | 2.374 | 1.67E-11 |
| KRAS | Activated | 2.243 | 1.83E-11 |
| CSF2 | Activated | 2.982 | 2.39E-11 |
| sodium arsenite | Activated | 2.032 | 3.74E-11 |
| RELA | Activated | 2.646 | 3.94E-11 |
| CSF1 | Activated | 2.605 | 5.57E-11 |
| arsenite | Activated | 2.93 | 6.25E-11 |
| deferoxamine | Activated | 2.587 | 7.83E-11 |
| NFKB1 | Activated | 2.933 | 8.75E-11 |
| HRAS | Activated | 2.636 | 9.94E-11 |
| TFRC | Inhibited | -2.848 | 1.29E-16 |
| PD98059 | Inhibited | -3.741 | 2.31E-15 |
| U0126 | Inhibited | -3.154 | 1.79E-13 |
| LY294002 | Inhibited | -2.875 | 2.35E-13 |
| SB203580 | Inhibited | -3.512 | 2.86E-11 |

**Supplementary Table 2:** IPA diseases & functions (all genes)

**Supplementary Table 2**

| Diseases or Functions Annotation | Predicted Activation State | Activation z-score | p-value |
|---|---|---|---|
| Cell movement | Increased | 2.613 | 2.18E-24 |
| Cell survival | Increased | 3.855 | 1.03E-23 |
| Migration of cells | Increased | 2.412 | 4.03E-22 |
| Cell viability | Increased | 3.856 | 1.15E-20 |
| Cancer | Increased | 2.633 | 4.53E-16 |
| Mammary tumor | Increased | 3.089 | 1.51E-12 |
| Cellular homeostasis | Increased | 2.572 | 6.7E-12 |
| Transmigration of cells | Increased | 2.039 | 7.69E-11 |
| Morbidity or mortality | Decreased | -2.675 | 6.86E-18 |
| Organismal death | Decreased | -2.704 | 7.02E-18 |

**Supplementary Table 3:** cyto PELP1 gene signature

**Supplementary Table 3**
**Cyto PELP1 Gene Signature**

| **Upregulated** | **Log2** | | |
|---|---|---|---|
| ADM | 1.10 | NA | NA |
| ANGPTL4 | 1.73 | -0.41 | NA |
| ASNS | 1.05 | 0.78 | 0.91 |
| BCL2 | 1.13 | 0.88 | 0.93 |
| BNIP3 | 1.19 | NA | 0.23 |
| BNIP3L | 1.02 | NA | 0.43 |
| CA9 | 2.30 | NA | NA |
| CA12 | 1.54 | 0.74 | 0.34 |
| EIF4EBP1 | 1.19 | 0.70 | 0.93 |
| ENO2 | 1.47 | 0.70 | 1.00 |
| EPAS1 | 1.22 | NA | NA |
| GHR | 1.12 | 0.90 | NA |
| INSIG2 | 1.13 | 0.66 | 0.96 |
| INSR | 1.02 | 0.88 | 0.98 |
| MED13L | 1.09 | 0.85 | 0.79 |
| MGST1 | 1.34 | NA | 0.95 |
| MT2A | 1.34 | NA | 0.95 |
| NDRG1 | 2.26 | NA | 0.58 |
| P4HA1 | 1.35 | 0.53 | 0.56 |
| PFKFB3 | 1.12 | NA | NA |
| PFKFB4 | 1.07 | -0.21 | -0.19 |
| PHGDH | 1.51 | 0.46 | 0.61 |
| PPFIA4 | 1.16 | NA | NA |
| RET | 1.01 | 0.70 | NA |
| SLC7A11 | 1.74 | 0.81 | 0.87 |
| SOX9 | 1.54 | 0.51 | NA |

| **Downregulated** | **Log2** | | |
|---|---|---|---|
| FSCN1 | -1.12 | -0.66 | -0.97 |
| HES6 | -1.19 | -0.92 | -0.70 |
| S100A8 | -1.06 | NA | -0.97 |
